# Sexually dimorphic architecture and function of a mechanosensory circuit in *C. elegans*

**DOI:** 10.1101/2022.02.18.481005

**Authors:** Hagar Setty, Yehuda Salzberg, Shadi Karimi, Elisheva Berent-Barzel, Michael Krieg, Meital Oren-Suissa

## Abstract

How sensory perception is processed by the two sexes of an organism is still only partially understood. Despite some evidence for sexual dimorphism in auditory and olfactory perception, whether touch is sensed in a dimorphic manner has not been addressed. Here we find that the neuronal circuit for tail mechanosensation in *C. elegans* is wired differently in the two sexes and employs a different combination of sex-shared sensory neurons and interneurons in each sex. Reverse genetic screens uncovered cell- and sex-specific functions of the alpha-tubulin *mec-12* and the sodium channel *tmc-1* in sensory neurons, and of the glutamate receptors *nmr-1* and *glr-1* in interneurons, revealing the underlying molecular mechanisms that mediate tail mechanosensation. Moreover, we show that only in males, the sex-shared interneuron AVG is strongly activated by tail mechanical stimulation, and accordingly is crucial for their behavioral response. Importantly, sex reversal experiments demonstrate that the sexual identity of AVG determines both the behavioral output of the mechanosensory response and the molecular pathways controlling it. Our results present for the first time extensive sexual dimorphism in a mechanosensory circuit at both the cellular and molecular levels.

## INTRODUCTION

In sexually reproducing species, evolutionary forces have shaped the nervous systems of the two sexes such that they feature overt sex-specific behaviors. These dimorphic behaviors can result from differences in sensory neuron perception, downstream neuronal processing, or both. Sexually dimorphic sensory perception has been observed in both vertebrate and invertebrate models. In *C. elegans*, the differential expression of a G-protein-coupled receptor (GPCR) in a sensory neuron determines the different behavioral outcome^1^. In frogs, sexually dimorphic auditory tuning has been documented^2–4^. In mice, sexually dimorphic olfactory perception^5,6^ and integration of sensory information were observed. The sexually dimorphic behavior in response to the pheromone ESP1, for example, was shown to be mediated through dimorphic processing in third- and fourth-order brain areas^7,8^. Integration downstream to the sensory level is also evident in aggressive behavior in *Drosophila* and nociceptive behavior in *C. elegans*, where sexually dimorphic processing by downstream interneurons regulates the dimorphic behavior^9,10^. These examples demonstrate that for different types of sensory modalities, be it olfactory, auditory or chemo-aversion, sex differences can originate from dimorphism in distinct neuronal layers.

The perception of mechanical forces, or mechanosensation, includes the perception of touch, hearing, proprioception and pain, all of which involve the transduction of mechanical forces into a cellular signal^11^. Numerous studies have focused on understanding the molecular and cellular pathways involved in mechanosensation in both mammalian and invertebrate systems^12–14^. Although recent evidence has provided insight into sex differences in mechanical nociception^15–19^, it is still unknown whether males and females sense innocuous touch differently.

*C. elegans* has been used extensively to explore the cells and molecules that govern mechanosensation, and specifically touch sensation^14,20^. Touch sensation in this species can generally be divided into two different modalities: gentle touch and harsh touch, each involving different cellular and molecular mechanisms^21–24^. The cellular mechanisms mediating harsh mechanical stimulation applied to the tail, or tail mechanosensation, have only recently begun to be examined. Studies focused on harsh-touch-induced tail mechanosensation discovered that the sensory neurons PHA, PHB and PHC and the interneurons PVC and DVA are required for tail mechanosensation in hermaphrodites^24,25^. These studies, however, did not identify the molecular mechanisms mediating touch sensitivity in these cells, nor did they address whether the same cells constitute or are part of the male tail mechanosensation circuit.

The neuronal connectome of both sexes in *C. elegans*^26–28^ indicates that many circuits, including those mediating mechanosensation, contain sexually dimorphic connections (Figure 1A). Here we report that while males and hermaphrodites respond similarly to harsh touch of the tail, they sense and integrate mechanosensory information differently, in terms of the neurons involved, the underlying circuits and the molecular components mediating and transducing touch. Our results establish the circuit architecture for tail mechanosensation in both sexes and reveal a novel role for the sex-shared interneuron AVG as an integrator of mechanosensory information. We further identify several key components of the molecular machinery controlling the activity of this circuit, and reveal the sex-specific functions of these molecules at both the sensory and interneuron levels. Our results demonstrate that the propagation of harsh-touch tail mechanosensory information is sexually dimorphic, and provide a unique example of how neuronal circuits evolved sex-specific features while maintaining the same sensory modality and its behavioral output.

**Figure 1.**
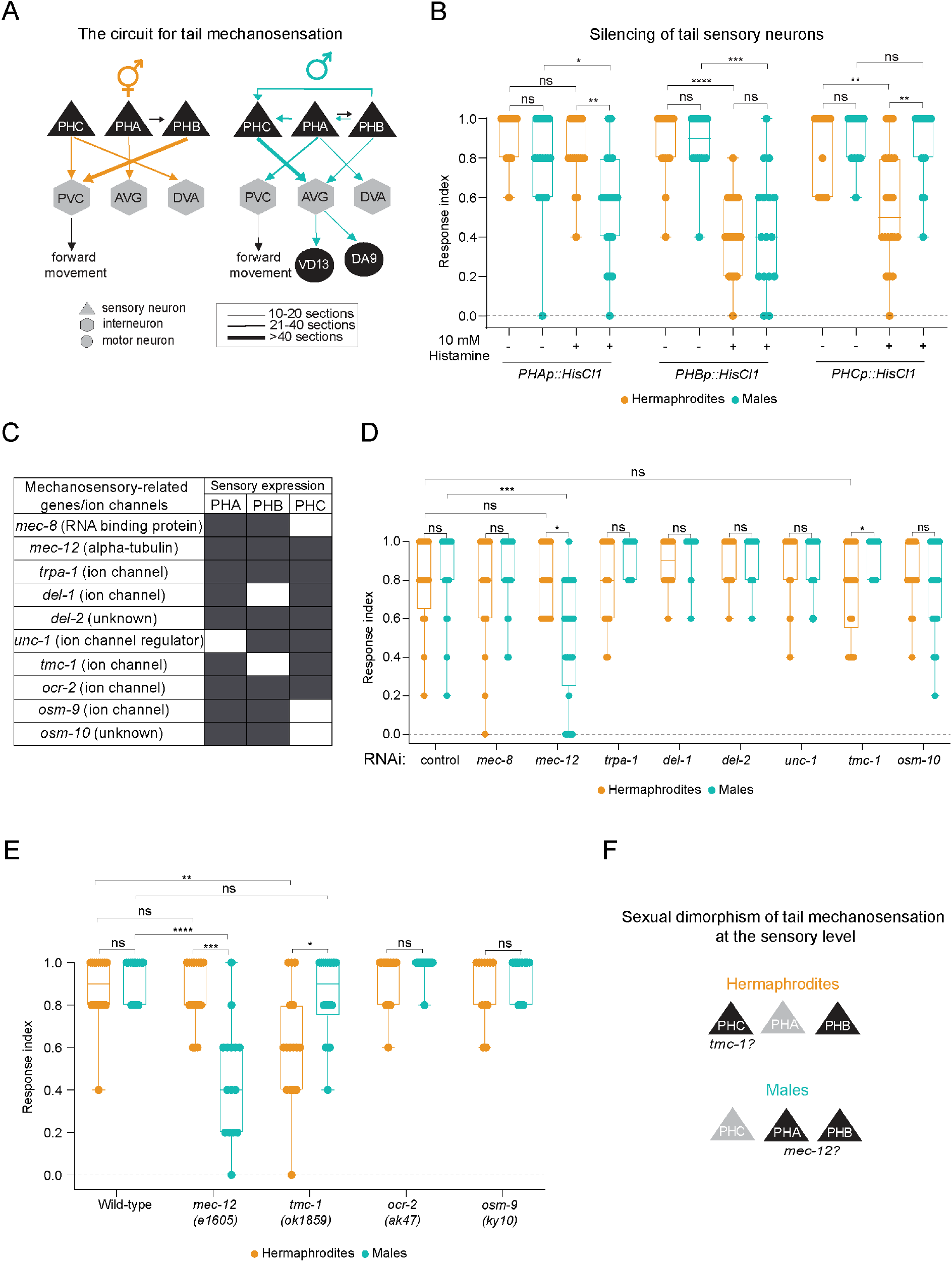
Sexually dimorphic perception of tail mechanosensation at the sensory level. (A) Predicted connectivity of the circuit for tail mechanosensation^26,28^. Chemical synapses between sensory (triangles), inter-(hexagons) and motor (circles) neurons are depicted as arrows. Thickness of arrows correlates with degree of connectivity (number of sections over which *en passant* synapses are observed). (B) Tail-touch responses of *PHAp::HisCl1-*, *PHBp::HisCl1-* and *PHCp::HisCl1*-expressing animals of both sexes that were tested either on histamine or control plates (see *Methods*). (C) Table listing the genes selected for reverse genetic screen and their expression pattern at the sensory neurons PHA, PHB and PHC. Dark boxes represent gene expression. (D) Tail-touch responses of RNAi-silenced candidate genes in both sexes. (E) Tail-touch responses of mutant strains for candidate genes in both sexes and in *him-5(e1490)* animals (see *Methods*). (F) Schematic of the sensory neurons which function in each sex, with the respective suggested sites of action of *tmc-1* and *mec-12*. The response index represents an average of the forward responses (scored as responded or not responded) in five assays for each animal. n = 11-21 worms per group. A Kruskal-Wallis test followed by Dunn’s multiple comparison test was performed for all comparisons, **** *p* < 0.0001, *** *p* < 0.001, ** *p* < 0.01, * *p* < 0.05, ns - non-significant. Orange-hermaphrodites, cyan-males.

## RESULTS

### Sensory level sexual dimorphism in the tail mechanosensation circuit

By combining data from the published connectome maps of both *C. elegans* sexes^26^ with behavioral data from previous studies^24,25^, we revealed that some of the sensory cells in the tail mechanosensation circuit are connected differently in the two sexes (Figure 1A). To explore the contribution of each sensory neuron to tail mechanosensation, we silenced individual neurons by cell-specific expression of the inhibitory *Drosophila* histamine-gated chloride channel (HisCl1)^29^ and then tested both sexes for tail-touch response (see *Methods*). Silencing the PHB neuron diminished the tail-touch response equally in both sexes, suggesting it has a sex-independent role in touch sensitivity (Figure 1B). However, silencing PHA and PHC revealed that PHC is necessary only in hermaphrodites (corroborating previous findings)^25^, while PHA is required only in males (Figure 1B). These results show that, at the sensory level, each sex utilizes a different combination of sensory cells in tail-touch perception.

We next asked whether the molecular mechanisms that govern tail mechanosensation at the sensory level are, too, sexually dimorphic. We carried out a reverse genetic screen targeting ion channels and other proteins previously suggested to be involved in mechanosensation that are known to be expressed in the tail sensory cells^21,30–36^ (Figure 1C). Assaying tail-touch responses of RNA interference (RNAi)-fed or mutant animals led to the identification of two genes whose silencing caused sex-specific defects in tail mechanosensation.

First, targeting *mec-12* by using both RNAi or a *mec-12* mutant, reduced tail mechanosensation only in males (Figure 1D-E). *mec-12* (expressed in PHA, PHB and PHC, Figure 1C) encodes an alpha-tubulin protein, is one of several genes required for touch receptor neuron (TRN) function in *C. elegans* and is specifically responsible for generating 15-protofilament microtubules in TRNs^37^.

Second, we found that RNAi of *tmc-1* elicited a reduced response only in hermaphrodites, and *tmc-1* mutant hermaphrodites exhibited a significantly reduced tail-touch response (Figure 1D-E). TMC-1 (expressed in PHA and PHC, Figure 1C) is a mechanosensitive sodium channel and an ortholog of the mammalian TMC proteins important for hair-cell mechanotransduction^31,38^. Taken together, our screen uncovered two molecules that play a role in mediating tail mechanosensation in a sex-dependent manner, possibly functioning through different types of sensory cells (Figure 1F). Overall, our findings demonstrate extensive cellular and molecular sexual dimorphism in mechanosensation at the sensory level.

### Cell- and sex-specific function of *mec-12* and *tmc-1* in tail mechanosensation

Having established a role for *mec-12* in mechanosensation in males, we turned to explore whether it functions sex-specifically through the phasmid neurons. Since PHA and PHB are required for tail mechanosensation in males (Figure 1B), we restored the expression of *mec-12* in mutant animals under the *che-12* promoter, which drives expression in ciliated amphid and phasmid neurons, including PHA and PHB^39^. We found that *mec-12* expression in ciliated neurons in *mec-12* mutant males is sufficient to rescue the tail-touch phenotype (Figure 2A), suggesting that *mec-12* functions in PHA/PHB. Although *mec-12* is required only in male PHA/PHB for tail mechanosensation, its expression in these cells was similar in the two sexes (Figure 2B-C).

**Figure 2.**
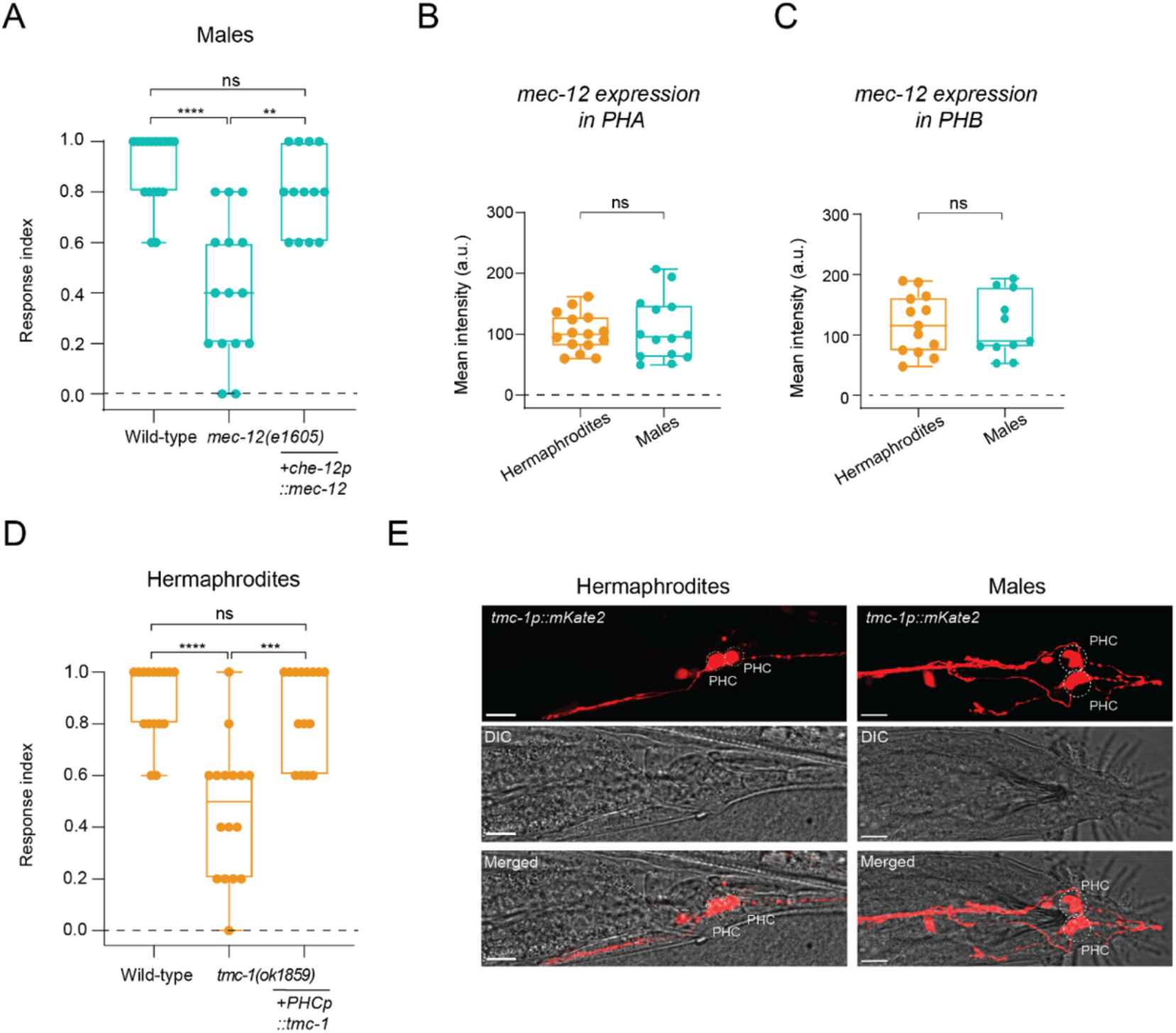
*mec-12* and *tmc-1* function cell- and sex-specifically in tail mechanosensation. (A) Tail-touch responses of wild-type, mec-12(e1605) and mec-12(e1605);che-12p::mec-12 males. n = 13-17 worms per group. (B-C) Quantification and comparison of *mec-12p::GFP* expression levels in PHA (B) and PHB (C). n = 11-14 worms per group. (D) Tail-touch responses of wild-type, *tmc-1(ok1859)* and *tmc-1(ok1859);PHCp::tmc-1* hermaphrodites. n = 15-17 worms per group. (E) Representative confocal micrographs of *tmc-1p::mKate2* in both sexes. Scale bars are 10 μm. PHC was identified using the *otIs520* transgene^25^. The response index represents an average of the forward responses (scored as responded or not responded) in five assays for each animal. For (B) and (C), we performed a Mann-Whitney test for each comparison. For (A) and (D), we performed a Kruskal-Wallis test followed by a Dunn’s multiple comparison test for all comparisons, **** *p* < 0.0001, *** *p* < 0.001, ** *p* < 0.01, ns - non-significant. Orange-hermaphrodites, cyan-males.

*tmc-1* has been shown to be expressed in PHA and PHC in hermaphrodites^33^ (Figure 1C). Since PHC is required for tail mechanosensation in hermaphrodites and not PHA (Figure 1B), we hypothesized that *tmc-1* mediates tail mechanosensation through PHC in hermaphrodites. Indeed, expressing *tmc-1* specifically in PHC rescued the defective tail-touch response of *tmc-1* mutant hermaphrodites (Figure 2D). This finding is supported by the evident expression of a *tmc-1::mKate2* transcriptional reporter in PHC in both sexes (Figure 2E). Taken together, our results demonstrate that not only do different sensory neurons participate in the processing of mechanosensory information in each sex, the molecules involved in this processing in the sensory neurons are, too, distinct between the sexes.

### The tail mechanosensation circuit is sexually dimorphic at the interneuron level

The predicted sexually dimorphic connectivity of the sensory neurons (Figure 1A) suggests dimorphic activities for the downstream interneurons. The sex-shared interneuron AVG is predicted to possess a striking dimorphic connectivity pattern according to the published connectomes, receiving more inputs in males compared to hermaphrodites (Figure S1A-B). Given that the sensory cells connected to AVG in males are the ones with a suggested role in mechanosensation, we speculated that AVG might be involved in tail mechanosensation. Therefore, we silenced AVG using a cell-specific driver (Figure S2A-B) and tested animals for tail-touch responses in both sexes. We found that silencing AVG elicits a sexually dimorphic effect on tail mechanosensation, reducing only the male response (Figure 3A), in agreement with AVG’s predicted dimorphic connectivity.

**Figure 3.**
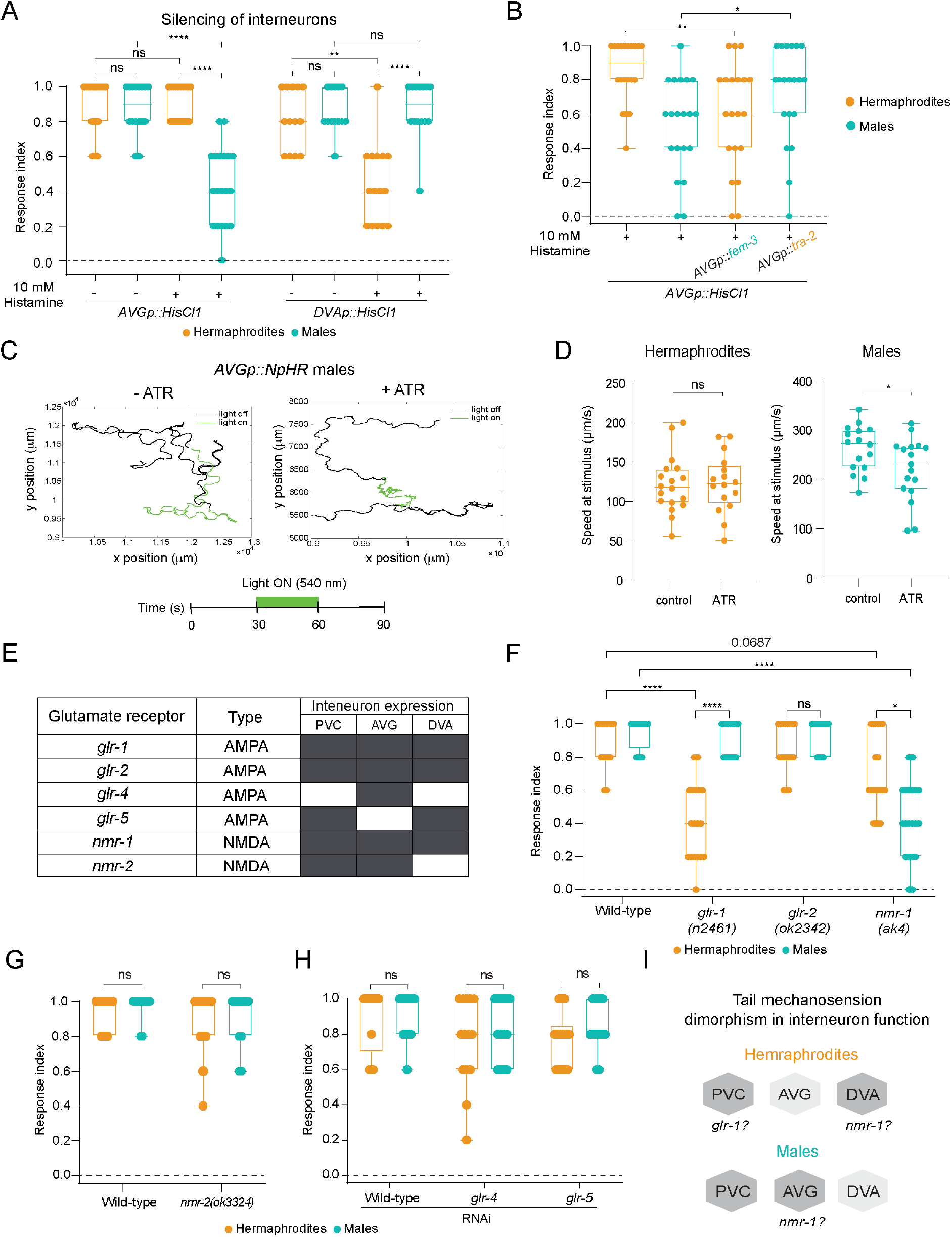
Sexually dimorphic integration of tail mechanosensation at the interneuron level. (A) Tail-touch responses of *AVGp::HisCl1-* and *DVAp::HisCl1*-expressing hermaphrodite and male worms that were tested either on histamine or control plates (see *Methods*). n = 14-21 worms per group. (B) Tail-touch responses of *AVGp::HisCl1* hermaphrodites (control and *AVGp::fem-3*) and males (control and *AVGp::tra-2*) that were tested on histamine plates. n = 20 worms per group. (C) Representative graph showing the head trajectory (head position) in a control male and ATR-supplemented male (upper panel). The schematic represents the timeline of the experimental setup (bottom panel). (D) Speed (μm/s) of hermaphrodite and male worms grown on control and ATR plates at the time of light projection. n = 16-18 worms per group. (E) Table listing the glutamate receptors selected for the reverse genetic screen, their expression pattern at the relevant interneurons and their predicted type. Dark boxes represent gene expression. (F-H) Tail-touch responses of gene candidates that were examined using mutant strains (F-G) or RNAi feeding (H) in both sexes. Each experiment was conducted with a control (*him-5(e1490)* (F)*, him-8(e1489)* (G) and *him-5(e1490)* fed with RNAi (H)). n = 12-20 worms per group. (I) Interneuron-level sexual dimorphism of tail mechanosensation with suggested mode of function for *nmr-1* and *glr-1*. The response index represents an average of the forward responses (scored as responded or not responded) in five assays for each animal. For (D), we performed a Mann-Whitney test for each comparison. For (A-B) and (F-H), we performed a Kruskal-Wallis test followed by a Dunn’s multiple comparison test for all comparisons, **** *p* < 0.0001, ** *p* < 0.01, * *p* < 0.05, ns - non-significant. Orange-hermaphrodites, cyan-males.

If AVG connectivity to the sensory neurons plays a significant role, rewiring AVG’s connections should affect tail-mechanosensation responses. Namely, adding connections in hermaphrodites would elicit an effect, and vice-versa for males. We therefore introduced a sex-determining factor specifically in AVG to switch its sexual identity and connectivity to that of the opposite sex^40–42^. We found that sex-reversing AVG is sufficient to convert the phenotype of the AVG-silenced tail-touch response: In hermaphrodites with a masculinized AVG, the tail-touch response was impaired when AVG was silenced compared to wild-type hermaphrodites, and in males with feminized AVG, the tail-touch response was rescued when AVG was silenced compared to wild-type males (Figure 3B). These results suggest that the sexual identity of AVG and consequently its wiring pattern, can shape tail mechanosensory behavior. Since the behavioral output of harsh touch applied to the tail is a forward movement, we asked whether optogenetic activation of AVG will result in forward movement only in males, as was shown for the sensory phasmid neurons in hermaphrodites^24^. Optogenetic activation of AVG did not affect the forward or total (forward+reverse) speed of the animals in both sexes (Figure S3). However, optogenetic inhibition of AVG reduced the total speed of males only (Figure 3C-D). Thus, AVG is required for locomotion in a sexually dimorphic manner.

We next tested whether the sex-shared interneurons DVA and PVC also have a dimorphic role in tail mechanosensation. As we were unable to generate a PVC-specific driver, in accordance with previous observations (Figure S4^43^), we focused on the role of DVA. A tail-touch assay on DVA-silenced animals revealed that only hermaphrodites are affected (Figure 3A, Figure S2C), in agreement with the predicted connectivity (Figure 1A). Taken together, our results uncover the sex-specific use of different interneurons in the circuit for tail mechanosensation, and assign a novel functional role for AVG in locomotion and mechanosensation.

Since we uncovered sexually dimorphic functions of the interneuron level in the circuit, we explored potential molecular mechanisms that might govern these differences. To this end, we screened a list of glutamate receptor genes expressed in the relevant interneurons (AVG, DVA and PVC^33,44^; Figure 3E), as the sensory neurons required for tail mechanosensation are glutamatergic^45^. We found that two glutamate receptors are required for tail mechanosensation in a sexually dimorphic manner: *glr-1* (AMPA type) is needed only in hermaphrodites, while *nmr-1* (NMDA type), which operates in both sexes, has a stronger effect in males (Figure 3F-H). Taken together, our data suggest that the dimorphic nature of the tail mechanosensation circuit spans beyond mere neuronal connectivity to include also different receptor dependency (Figure 3I).

### NMDA receptor *nmr-1* is required specifically in AVG to mediate tail mechanosensation in males

We next sought to assess the cell-autonomous role of *nmr-1* and *glr-1* in tail mechanosensation in AVG. To do so, we overexpressed *nmr-1* and *glr-1* specifically in AVG and determined the tail-touch response in the respective mutants. We found that *nmr-1* re-expression in AVG, but not *glr-1* re-expression, rescues the defective tail-touch phenotype of mutant males and not of hermaphrodites (Figure 4A, Figure S5A), suggesting that *nmr-1* functions cell autonomously in male AVG for this purpose. In line with this observation, the expression of *nmr-1* fosmid in AVG was higher in males (Figure 4B-C). *glr-1* expression pattern was observed in AVG in both sexes but at higher levels in males (Figure S5B-C), suggesting it mediates a different and possibly dimorphic function in AVG.

**Figure 4.**
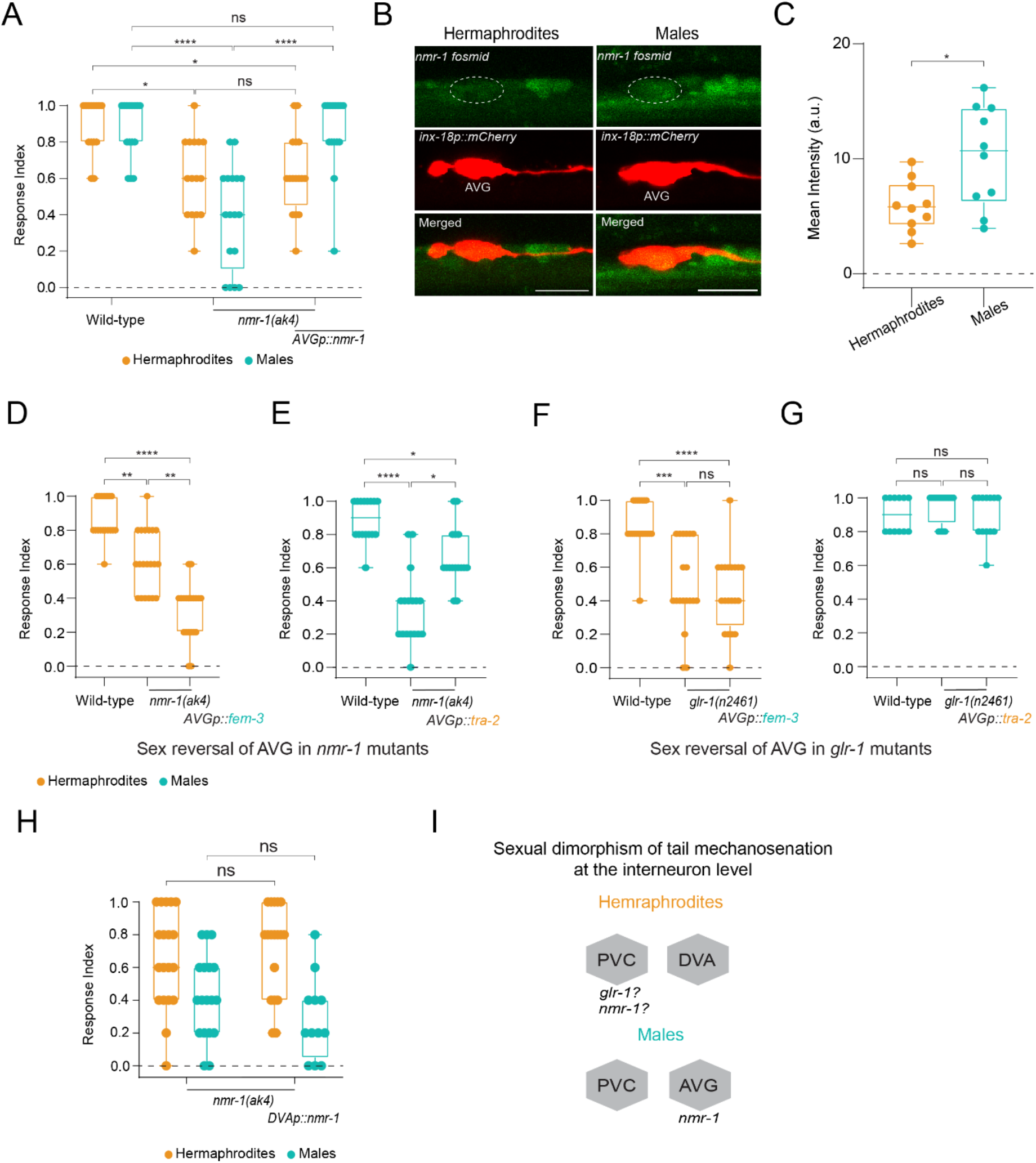
Cell-autonomous and sex-specific role for *nmr-1* in tail mechanosensation. (A) Tail-touch responses of wild-type, *nmr-1(ak4)* and *nmr-1(ak4);AVGp::nmr-1* hermaphrodites and males. n = 14-17 worms per group. (B) Representative confocal micrographs of *nmr-1::GFP* fosmid in the AVG interneuron, identified by the expression of mCherry in both sexes. Scale bar is 10 μm. (C) Quantification of (B). n = 10-11 worms per group. (D,F) Tail-touch responses of hermaphrodites with AVG masculinized on a *nmr-1(ak4)* mutant background (D) and *glr-1(n2461)* mutant background (F) with respective controls. n = 18-20 worms per group. (E,G) Tail-touch responses of males with AVG feminized on a *nmr-1(ak4)* mutant background (E) and *glr-1(n2461)* mutant background (G) with respective controls. n = 12-17 worms per group. (H) Tail-touch responses of wild-type, *nmr-1(ak4)* and *nmr-1(ak4);DVAp::nmr-1* hermaphrodites and males. n = 12-19 worms per group. (I) Receptor site-of-action for tail mechanosensation. The response index represents an average of the forward responses (scored as responded or not responded) in five assays for each animal. For (C), we performed a Mann-Whitney test. For (A) and (D-H), we performed a Kruskal-Wallis test followed by a Dunn’s multiple comparison test for all comparisons, **** *p* < 0.0001, *** *p* < 0.001, ** *p* < 0.01, * *p* < 0.05, ns - non-significant. Orange-hermaphrodites, cyan-males.

Since we observed that the sexual identity of AVG is sufficient to determine the behavioral outcome of the circuit when AVG is silenced (Figure 3B), we asked whether this is also true in *nmr-1* or *glr-1* mutant animals. Sex-reversing AVG in *nmr-1* mutant animals switched the tail-touch phenotype to that of the opposite sex, i.e., it reduced the tail-touch response in hermaphrodites with masculinized AVG and enhanced it in males with feminized AVG (Figure 4D-E). This was not evident in *glr-1* mutant animals, as sex-reversal of AVG had no effect in both sexes (Figure 4F-G). These results corroborate the observations of the cell-specific rescue experiments, and point to a cell-autonomous role for *nmr-1* in AVG in males that mediates tail mechanosensation. Importantly, the sexual identity of AVG not only dictates the connectivity pattern^40^, but also the cell-autonomous molecular pathway that mediates the behavior.

Our results also show a slight defect in tail mechanosensation in *nmr-1* mutant hermaphrodites (Figure 4A). This led us to search for the interneuron in hermaphrodites in which *nmr-1* mediates tail mechanosensation. Since DVA is required for tail-touch response only in hermaphrodites (Figure 3A), and *nmr-1* is known to be expressed in hermaphrodites in DVA (Figure 3E), we checked whether *nmr-1* functions through DVA to mediate tail mechanosensation. Re-expressing *nmr-1* in DVA did not rescue the response of *nmr-1* mutant hermaphrodites (Figure 4H), suggesting *nmr-1* might be required in a different interneuron, such as PVC, for tail mechanosensation (Figure 4I).

### Mechanical stimulation of the tail elicits a sexually dimorphic neuronal response in AVG

Since AVG has a role in integrating mechanical information specifically in males, we asked whether it is activated in response to the application of mechanical force to the tail, and whether such activation occurs sex-specifically. We recorded the calcium traces of AVG in both sexes in response to tail mechanical stimulation using a microfluidic device that was adjusted to fit the male body^46^ (Figure S6). Similar to our previous observation in touch receptor neurons^46^, we found that AVG exhibits blue-light-evoked Ca^2+^ transients even in the absence of a mechanical stimulus, suggesting a previously uncharacterized stimulatory effect of LITE-1 on AVG (Figure S7A-D). We therefore measured the mechanosensitive activity of AVG under a *lite-1* mutant background. Three consecutive tail mechanical stimulations, but not posterior stimulations (mock), elicited neuronal responses in AVG (Figure 5A-C). Importantly, these responses were sexually dimorphic, being significantly lower in hermaphrodites compared to males (Figure 5A-B; Figure S8A). These findings support our behavioral results, and further indicate that AVG integrates mechanosensory information in a dimorphic manner. In line with the cell-autonomous role we uncovered for *nmr-1* in AVG in tail mechanosensation, we found that AVG responses to mechanical stimulations were reduced in *nmr-1* mutant males compared to controls (Figure 5D, Figure S8B).

**Figure 5.**
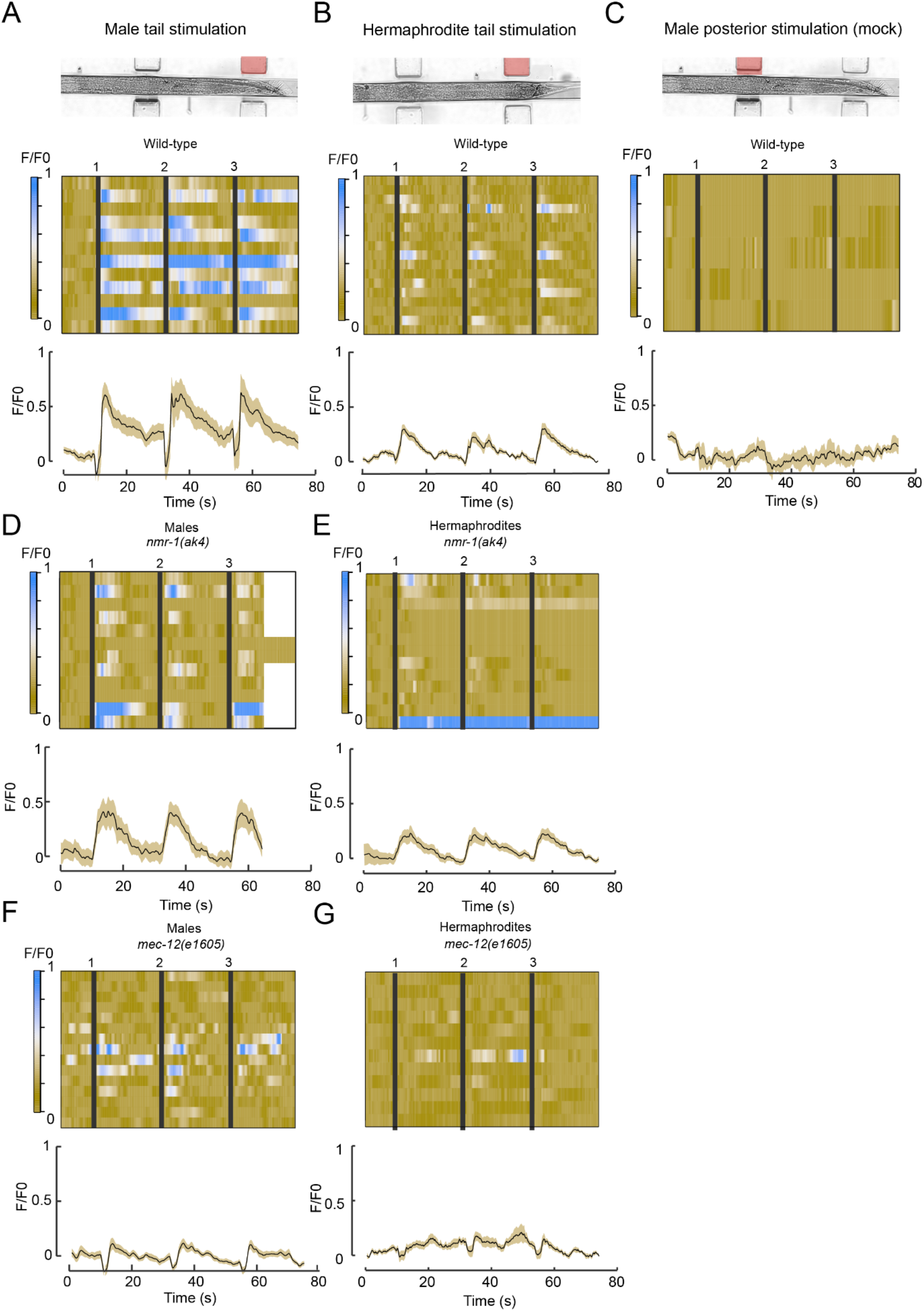
Sexually dimorphic responses of AVG to mechanical stimulation. (A-C) AVG GCaMP6s calcium responses of males and hermaphrodites to three consecutive tail mechanical stimulations (A-B) and of males to three consecutive posterior mechanical stimulations (C). Stacked kymographs represent the GCaMP intensity vs. time of individual recordings. Graphs represent average and SD traces of AVG calcium responses. Black vertical lines represent the time when a stimulus was applied. n = 6-17 animals per group. (D, E) AVG GCaMP6s calcium responses of *nmr-1(ak4)* mutant males (D) and hermaphrodites (E) to three consecutive tail mechanical stimulations. n = 13-14 animals per group. (F, G) AVG GCaMP6s calcium responses of *mec-12(e1605)* mutant males (F) and hermaphrodites (G) to three consecutive tail mechanical stimulations. n = 12-28 animals per group. Each 3.5 bar stimulus was applied for two seconds (see *Methods*). Full statistical analysis can be found in Figure S8.

We next asked how the sensory processing of mechanical stimulation is translated at the interneuron level. To explore this issue, we recorded the calcium traces of AVG in response to tail mechanical stimulation in *mec-12* mutant males, where sensory processing of mechanical stimulation is compromised (Figure 2). Interestingly, *mec-12* mutant males showed significantly lower AVG responses compared to wild-type (Figure 5F, Figure S8C). This result indicates that proper sensory perception of mechanical stimulation through *mec-12* is critical for the integration at the interneuron level. We also observed lower AVG responses both in *nmr-1* and *mec-12* mutant hermaphrodites, suggesting a role for the two genes in AVG integration of mechanical stimulation in hermaphrodites as well (Figure 5E, G). Taken together, AVG integrates tail mechanosensation in a sexually dimorphic manner, and requires *mec-12-*dependent inputs and *nmr-1* for this purpose.

### Mating behavior is compromised in *nmr-1* mutant males

The male tail bears the copulatory apparatus and contains specialized sensory structures required for mating^47^. We thus wondered whether the male-specific use of particular neurons and genes for tail mechanosensation may reflect a broader role they have in the mating circuit. To test this, we performed mating assays on mutants for the male-specific genes we discovered, *mec-12* and *nmr-1*. While *mec-12* mutant males did not show any defects in mating, *nmr-1* mutant males were indeed defective in their ability to locate the hermaphrodite vulva, a crucial step in the mating sequence. *nmr-1* mutant males also displayed slight defects in their response to contact with hermaphrodites (Figure 6A-C). These findings suggest that at least some of the genes and mechanisms that mediate tail mechanosensation in males may have been “hijacked” from or by the mating circuit, where they serve additional functions.

**Figure 6.**
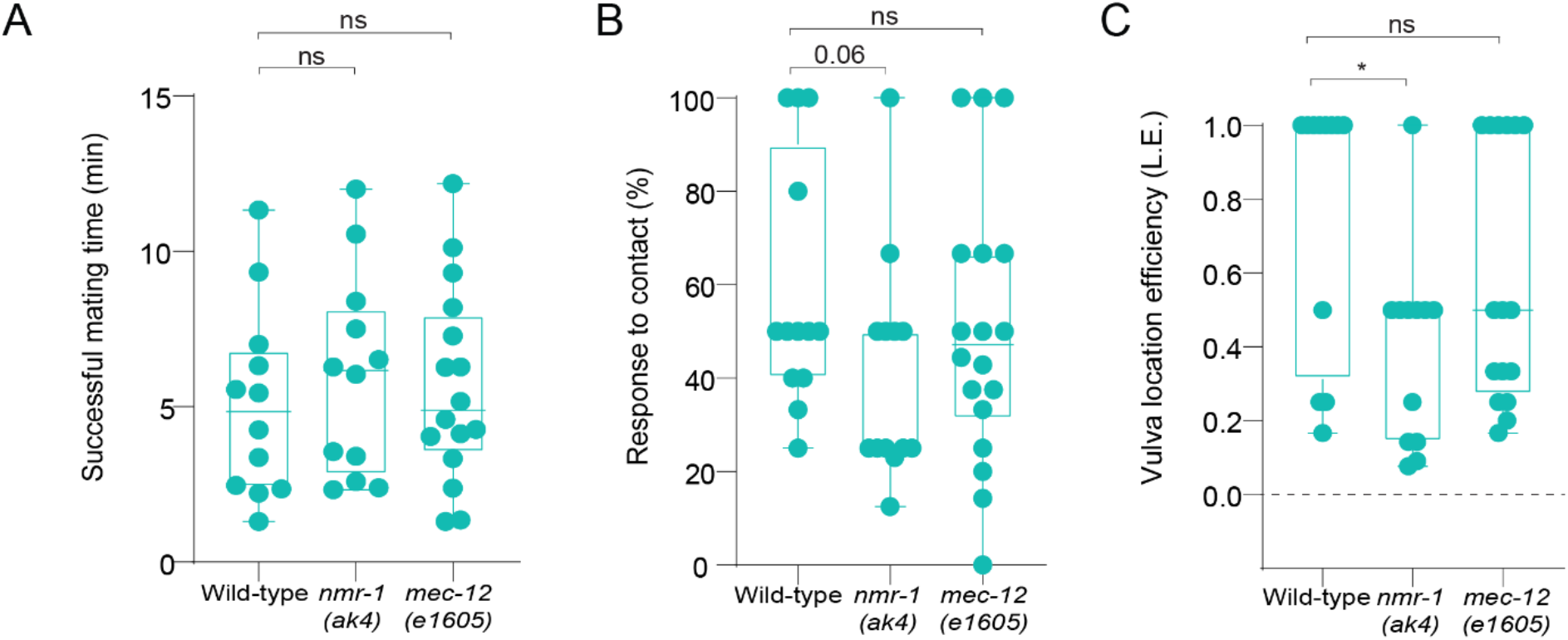
*nmr-1* is required for male mating behavior. Quantification of time until successful mating (A), response to contact (B) and vulva location efficiency (L.E., see *Methods*) (C) in wild-type, *nmr-1(ak4)* and *mec-12 (e1605)* males. n = 12-18 worms per group. We performed a Mann-Whitney test for each comparison. * *p* < 0.05, ns - non-significant.

**Figure 7.**
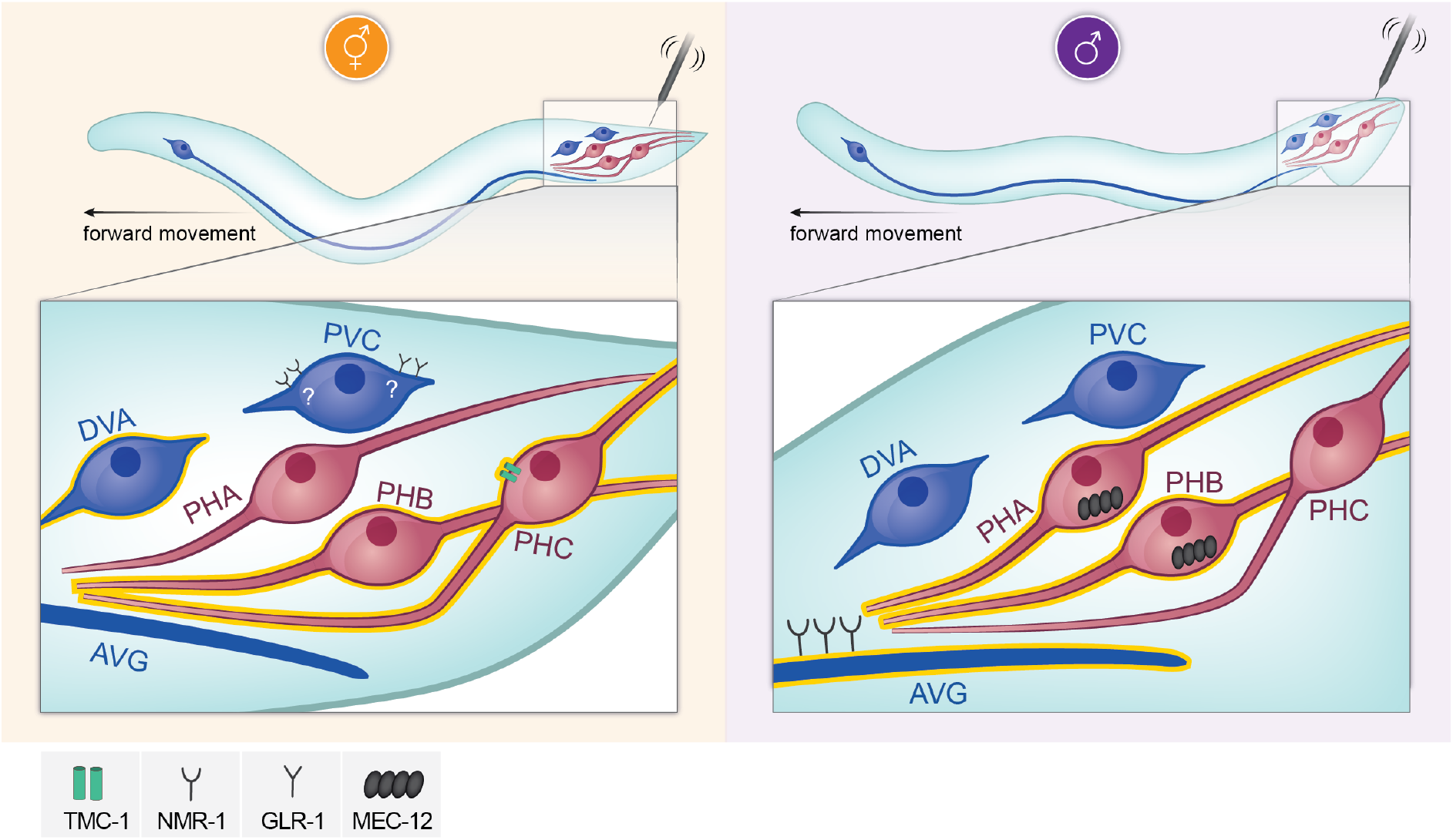
Sexually dimorphic cellular and molecular mechanisms controlling tail mechanosensation. A model depicting the cellular and molecular elements mediating tail mechanosensation in each sex. Neurons that function in each sex are highlighted in yellow.

## DISCUSSION

In this study, we present a unique example for a simple sensory circuit that underwent extensive sexually dimorphic reorganization during evolution, engaging different combinations of cells, connections and molecular pathways in the two sexes to mediate the same sensory modality, i.e., tail mechanosensation. While very few studies have analyzed the dimorphic properties of sensory circuits in detail in vertebrates, several examples exist in *C. elegans* for either cellular or molecular sensory dimorphism. For example, hermaphrodites detect food-related olfactory cues better than males due to enhanced expression of the odorant receptor ODR-10 in the AWA neuron^1^. Conversely, in the anterior nociceptive circuit, sensory detection of aversive cues seems identical in the two sexes, but the downstream connectivity to interneurons is highly dimorphic^10^. These topographical differences drive sexually dimorphic behavioral responses to nociceptive cues. Notably, in these examples, the dimorphic properties of the circuit lead to dimorphic behavioral outputs, whereas in the circuit we studied here, wild-type males and hermaphrodites respond to the stimulus in the same manner, suggesting that this modality was important enough for both sexes to withstand the evolutionary pressures that changed its constituents. One such pressure on the male circuit was the need to integrate the mating circuit into the shared nervous system, as suggested by the involvement of the same cells (e.g., AVG and PHB in this study, and see also^40^) and genes (*nmr-1* in this study) in tail mechanosensation and mating behaviors.

We found four genes that function sex-specifically in the circuit: *mec-12* and *tmc-1* in sensory cells, and *glr-1* and *nmr-1* in interneurons. Previous reports indicate that the alpha-tubulin MEC-12 is required with the beta-tubulin MEC-7 for 15-protofilament microtubule assembly in touch receptor neurons (TRNs), responsible for the transduction of gentle touch^37,48^. Our data suggests that MEC-12 functions through PHA/PHB to mediate tail mechanosensation in males, but whether this function requires MEC-7 remains to be determined. The involvement of PHA and PHB, two ciliated neurons, in tail mechanosensation in males and not the non-ciliated PHC suggests that the defects in *mec-12* mutant males are likely due to aberrant cilia formation in these neurons. It appears that MEC-12, a molecule required for gentle touch^21^, is also required for harsh touch, suggesting some overlap between the molecular mechanisms mediating the two modalities, as was previously shown for MEC-10 and MEC-3^22,23^.

We also found that the sodium-sensitive channel TMC-1^31^ is a hermaphrodite-specific mediator of tail mechanosensation. It was previously shown that TMC-1 is required in *C. elegans* for salt sensation, avoidance of noxious alkaline environments, egg laying, gentle-nose touch response, and the inhibition of egg-laying in response to a harsh mechanical stimulus^31,38,49–51^. Together with our findings, TMC-1 appears to function as a polymodal ion channel, enabling the processing of different types of information. In *Drosophila*, *tmc* is required for mechanosensitive proprioception^52^, whereas in mice, *Tmc1* is expressed in cochlear hair cells and is required for proper mechanotransduction^53–56^. In both mice and humans, dominant and recessive mutations in *Tmc1* leading to deafness have been identified^53,56^. Sex differences in hearing loss have been documented in mice^57–60^, raising the possibility that sex-specific mechanisms involving *Tmc1* may also exist in vertebrates. While a possible dimorphic role for *Tmc1* in higher organisms is yet to be discovered, our results highlight the importance of research on sex-differences for future sex-specific therapeutic approaches.

Our previous and current work together show that AVG is a key dimorphic interneuron, responding differently in males and hermaphrodites to at least two separate modalities, tail mechanosensation and nociception^61^. The response of AVG to mechanical stimulation is achieved only when the tail is stimulated but not when other posterior body areas are touched, emphasizing that the input arrives from the tail sensory neurons and that the response to a harsh touch in the tail and the posterior body are distinct modalities involving different cellular and molecular mechanisms^24^. Blue-light-evoked activity in AVG independently of mechanical stimulation suggests that LITE-1 has a role in or upstream to AVG. Supporting this possibility are the findings that AVG is the neuron with the strongest expression of *lite-1*^33^ and that PHA requires LITE-1 for H2O2 sensation^62^.

We expose the intricate use of sex-shared interneurons as dimorphic hubs for different circuits. For example, the AVG interneuron is necessary only in males both for tail-touch and for locomotion speed, whereas the DVA interneuron is necessary for tail-touch only in hermaphrodites and for locomotion speed only in males^63^. Interestingly, calcium recordings of AVG following tail touch reveal a response in hermaphrodites that does not translate into a behavioral output, suggesting a threshold mechanism in AVG.

The published connectivity maps suggest that PVC probably plays a critical role in the integration of mechanosensation in both sexes, together with AVG in males and DVA in hermaphrodites. Therefore, a combinatorial system of interneurons seems to be at play to mediate multiple different circuits in each sex.

Despite the superficial similarity in the response of both sexes to touch, we uncover here that the propagation of mechanosensory information is dramatically different, providing the first evidence for a sexually dimorphic integration of touch signals. Our results contribute to the understanding of how sexually dimorphic circuits function to provide the organism with both the sensation of the environment and the behaviors that ensure its fitness.

## ACKNOWLEDGEMENTS

We thank members of the Oren-Suissa lab for their critical insights regarding the manuscript. The plasmids to amplify Chr2 and NpHR were a generous gift from Alon Zaslaver. The primer sequences to generate pHS11 were a kind recommendation of Carmine Puckett Robinson. The pEK158 plasmid and the AQ4330 strain were a kind gift from Eva Kaulich and Bill Schafer. Some strains were provided by the CGC, which is funded by the NIH Office of Research Infrastructure Programs (P40 OD010440).

This work was supported by the European Research Council ERC-2019-STG 850784 (MOS), Israel Science Foundation grant 961/21 (MOS). MOS is grateful to the Azrieli Foundation for the award of an Azrieli Fellowship, and is the incumbent of the Jenna and Julia Birnbach Family Career Development Chair. MK acknowledges financial support from the ERC (MechanoSystems, 715243), HFSP (CDA00023/2018), Spanish Ministry of Economy and Competitiveness through the Plan Nacional (PGC2018-097882-A-I00), FEDER (EQC2018-005048-P), “Severo Ochoa” program for Centres of Excellence in R&D (CEX2019-000910-S; RYC-2016-21062), Fundació Privada Cellex, Fundació Mir-Puig, and from Generalitat de Catalunya through the CERCA and Research program (2017 SGR 1012).

## AUTHOR CONTRIBUTIONS

HS, YS conducted and analyzed the experiments, SK conducted and analyzed the calcium imaging experiments, EBB contributed to behavioral experiments (“tail-touch assays”). MK and MOS supervised and designed the experiments. HS, YS and MOS wrote the paper.

## METHODS

### C. elegans strains

Wild-type strains were *C. elegans* variety Bristol, strain N2. *him-5(e1490)* or *him-8(e1489)* were treated as wild-type controls for strains with these alleles in their background. Worms were maintained according to standard methods^64^. Worms were grown at 20°C on nematode growth media (NGM) plates seeded with bacteria (*E. coli* OP50) as a food source. All the transgenic animals used in this study are listed in Supplementary Table 1, ordered by figures.

### Histamine-induced silencing

Histamine plates were prepared as previously described^29^. NGM-histamine (10 mM) and control plates were stored at 4°C for no longer than 2 months. Histamine plates were tested using worms that carry a transgene with a pan-neuronal HisCl1 (*tag-168::HisCl1::SL2::GFP*)^29^. After a few minutes on histamine plates, these worms were paralyzed completely, validating the potency of the histamine plates.

### Tail-touch assay

The assay was based as described^24^. In brief, L4 animals were isolated the day before the experiment and stored at 20°C overnight. On the day of the experiment, single 1-day adult animals were transferred into NGM plates freshly seeded with 30 μl OP50. After 30 minutes of habituation, animals were tested by applying a touch to the tail with a flattened platinum wire pick. Touch was applied to worms that did not move or moved very slowly. Each worm was tested five times with intervals of at least 10 seconds between each trial. The tested worm was given a score of one if it moved forward in response to the touch, and zero if it did not move forward. The response index was then calculated as the average of the forward responses. For assays using histamine-gated chloride channels, animals were grown on histamine plates or control plates seeded with 200-300 μl OP50 overnight. On the day of the experiment, single 1-day adult animals were assayed on histamine plates or control plates seeded with 30 μl OP50. For assays using RNAi, *him-5(e1490)* animals were grown on RNAi plates or control plates seeded with 200 μl of the RNAi bacteria.

### Molecular cloning

To generate the *mec-12* rescue construct pHS16 (*che-12p::mec-12*), *mec-12* cDNA was amplified from a N2 mixed-stage cDNA library and cloned by Gibson assembly^65^ into a che-12p plasmid backbone.

To generate the optogenetics plasmids (pHS5 - *inx-18p::NpHR::mCherry* and pHS6 - *inx-18p::Chr2::mCherry),* NpHR and Chr2 were amplified from plasmids provided by Alon Zaslaver and cloned into pMO46^40^.

To generate the *nmr-1* rescue constructs pHS14 (*inx-18p::nmr-1* cDNA) and pYS53 (*WT300::pes-10::nmr-1* cDNA), *nmr-1* cDNA was amplified from a N2 mixed-stage cDNA library and cloned by Gibson assembly into an *inx-18p* plasmid backbone, and pHS11 (*WT300::pes-10::mCherry*, swapping the mCherry fragment), respectively. WT300 was amplified from genomic DNA^66^.

To generate pYS52 (*WT300::pes-10::HisCl1::SL2::GFP*), HisCl1::SL2::GFP was amplified from pMO46^40^ and cloned into pHS11 (WT300::pes-10::mCherry, swapping the mCherry fragment) using Gibson assembly.

To express *tmc-1* cDNA specifically in PHC, *tmc-1* cDNA was amplified from pEK158 (kind gift of Eva Kaulich and Bill Schafer) and fused to a plasmid backbone containing the *eat-4p11* promoter by Gibson assembly, creating pYS54.

### RNAi

RNA interference was performed using the feeding method^67^. L4 hermaphrodites were fed HT115 bacteria carrying dsRNA for the relevant genes or a control empty vector, and their progeny were assayed for a tail-touch response at 1-day adult. *pos-1* RNAi served as a positive control for RNAi efficiency. A fresh *pos-1* control was prepared for each RNAi experiment.

### DiD staining

Worms were washed with M9 buffer and incubated overnight in 1 ml M9 and 5 μl DiD dye (Vybrant™ DiD Cell-Labeling Solution, ThermoFisher) at ~10-20 rpm. The worms were then centrifuged and transferred to a fresh plate.

### Microscopy

Animals were mounted on a 5 % agarose pad on a glass slide, on a drop of M9 containing 100-200 mM sodium azide (NaN3), which served as an anesthetic. A Zeiss LSM 880 confocal microscope was used with 63x magnification. For *nmr-1* fosmid, AVG was identified using *mCherry* expression, and the z-plane with the strongest signal was chosen to measure the fluorescence intensity using ImageJ version 1.52p. Regions of interest (ROIs) were selected manually with an auto local threshold using the Bernsen method. For *mec-12p::gfp* quantification in PHA/PHB, cell identification was done using the DiD stain, and ROIs were selected manually. Figures were prepared using Adobe Illustrator v24.0.

### Optogenetics

For optogenetic inhibition or activation, we used worms expressing NpHR or Chr2, respectively, only in AVG (*inx-18p::NpHR::mCherry/inx-18p::Chr2::mCherry*). L4 worms were picked a day before the experiment and separated into hermaphrodite and male control and experiment groups. They were transferred to newly seeded plates with 300 μl OP50 that was concentrated 1:10. ATR (all-trans-retinal) was added only to the experimental groups’ plates, to a final concentration of 100 μM. As ATR is sensitive to light, all plates were handled in the dark. Tracking and optogenetics was done on unseeded NGM plates. On the day of the experiment, the plates were seeded with 30 μl OP50, and ATR was added to the experimental group’s plates. To keep the worms in the camera field of view, a plastic ring was placed on the agar and a single worm was placed inside the ring. After 10 minutes of habituation, the worm was tracked for 90 seconds, with 30 seconds of 540 nm LED activation initiated 30 seconds after tracking commencement. The LED intensity was 0.167 mW/mm2. Speed measurements were extracted for the different time periods from WormLab (MBF Bio-science^68^) using “label-analysis”. The head position trajectory quadrants were also extracted from WormLab, and inserted into a graph using MATLAB.

For optogenetic activation, 3-5 worms were placed inside a plastic ring on the agar each time. After 10 minutes of habituation, the worms were tracked for 60 seconds, with a 2-second blue-light stimulus introduced every 10 seconds (5 stimuli in total). The LED intensity was 1.47 mW/mm2. Speed measurements were extracted for all the time points from WormLab. For each stimulus, the mean total speed was calculated before the stimulus (3 seconds) and after the stimulus (5 seconds). Calculations were done using MATLAB.

### Calcium imaging of animals in the microfluidic chip

#### Mold design and preparation

Microfluidic devices are based on the body wall touch device by^46^. Subtle modification in the trapping channel (width and height have been adjusted to 30 μm) were applied to fit male animals, which are skinnier than hermaphrodites, and the devices were cast with a PDMS base polymer/curing agent ratio to allow for large deflection of the diaphragm. The male body wall touch design, drawn with AutoCAD 2021, is illustrated in Figure S6A; its file is available to download in the Supplementary Material. Figure S6B shows how perfectly the modified trapping channel fits the males. To further enable a larger deflection, the size of the diaphragm was decreased to 10 microns. The deflection of the actuator under different pressure levels is presented in Figure S6C; the graph highlights the consistency of the numerical results with the experimental data.

The mold was prepared using a standard soft-lithography technique in a class ISO2 cleanroom^69^. In short, 4-inch silicon wafers were cleaned in Piranha and dehydrated by baking at 150°C. The dried, cleaned wafers were spin-coated with 1 ml/inch SU8-50 (MicroChem, Newton Massachusetts, USA) to obtain a 30-micron height (500 RPM (100 RPM/s acceleration for 15 seconds)) and then ramped to a final speed 3300 RPM for 45 seconds (300 RPM/s acceleration). The SU8 layer was pre-baked at 65°C for 5 minutes and then at 95°C for 15 minutes. The device design was ‘printed’ with a UV laser onto the SU8 substrate with a mask-less aligner (MLA, Heidelberg Germany). Once the printing process was finished, the wafer was post-baked for 1 minute at 65°C and then for 4 minutes at 95°C. The exposed SU8 was developed for 6 minutes in an SU8 developer (MicroChemicals) and rinsed with propanol, to remove unexposed resin. Thereafter, the wafer was hard-baked for 2h at 135°C and used for replica-molding PDMS devices. First, the surface of the structured wafer was vapor-phase silanized with Chlorodimethylsilane (Sigma-Aldrich, Missouri, United States) to prevent adhesion of the PDMS to the substrate and facilitate lift-off of the PDMS during the peeling process. The PDMS base polymer and curing agent were mixed at a ratio of 15:1 and approximately 40 mL per wafer were prepared and degassed before pouring to prevent bubbles. The degassed PDMS was poured into the mold and cured in the oven at 85°C for two hours. After lift-off and trimming the devices to fit onto standard #1.5 cover glasses, all inlets and outlets were punched with a biopsy punch (0.75 mm), which served as a receptacle for the connection tubes. Finally, to bind the PDMS with the cleaned cover slip, the surface of the coverslip was activated by a plasma treatment (Plasma Asher PVA TePla 300), and to increase the bonding quality, the bonded chip was placed on a hotplate (120°C) for 10 minutes. The punch holes were plugged with SC22/15-gauge metal tubes connected to 22-gauge catheter tubing.

#### Animal loading, stimulation and fluorescence microscopy

Animals were placed in the device exactly as described and shown in^70^. In brief, 3-4 synchronized day-one adult animals (male or hermaphrodites) were picked from an NGM plate containing OP50 bacteria and transferred into a 15 μL droplet of M9 buffer. Using a stereo dissecting scope (Leica S9), the animals were aspirated into a SC22/15-gauge metal tube (Phymep) connected to a 3 mL syringe (Henke Sass Wolf) with a PE tube (McMaster-Carr) pre-filled with M9 buffer. The loading tube was inserted into the inlet port of the device, while gentle pressure to the plunger of the syringe released the animals into the loading chamber. The microfluidic chip was then transferred to a compound fluorescence microscope (Leica DMi8), and with the 25X 0.95 water immersion lens, the animal was positioned such that the tail aligned with an actuator (Fig. 5A), ready to accept a mechanical stimulus. Once positioned properly, AVG was brought into focus and the stimulation protocol was started. GCaMP in AVG was excited with a cyan LED of a Lumencor SpectraX light engine (470 nm @ 2% transmission). Emission was collected through a 515/15nm emission filter (Semrock) and videos were captured at a 10 Hz acquisition rate (85 ms exposure time) with a Hamamatsu Orca Flash 4.3 for 65 or 75 seconds, using HCImage software. A master-pulse was used to synchronize the camera acquisition with the light exposure. Likewise, the camera SMA trigger was connected to the OB1 pressure controller and used to synchronize the image acquisition with the pressure protocol through the ElveFlow sequencer software prior to the imaging routine, ensuring precise and repetitive timing of the pressure protocol. The sequence consisted of 100 pre-stimulus frames and three consecutive mechanical stimuli, each separated by 20 seconds. The stimulus profile consisted of a 2-second 250 kPa pressure step, overlayed with a 150 kPa oscillation, resulting in a maximum deflection of 9 μm (Supplementary Figure S6C).

#### Analysis

Image processing and analysis were performed as described in^19^. To extract GCaMP intensity over time, images were preprocessed in ImageJ and then imported into python to extract signal intensity using in-house procedures written in Python or IgroPro. In short, the image sequence was cropped to a small area surrounding the cell body of the neuron of interest and a Gaussian filter was applied. All pixel values in the ROI were ranked and the background intensity was extracted from the 0-10th percentile and the GCaMP signal was extracted from the 90-100th percentile. After background subtraction, signal intensity was normalized to the first 100 frames (before the mechanical stimulus was applied).

To calculate a time-varying p-value comparing two datasets derived from different conditions, we first calculated the average calcium response after performing a non-linear baseline subtraction (bleach correction, drift) of each individual recording, taking advantage of an iterative algorithm to suppress the baseline by means in local windows Liland, MethodsX, 2015. To minimize spurious fluctuations in p-values due to imaging noise, a smoothing kernel based on the 2nd derivative penalty, known as a Whittaker smoother^71^, was applied. The *t*-test statistics was then used to calculate the *p*-value considering the degrees of freedom, *df=N1+N2-2*; the null hypothesis was rejected if the absolute value of *t* was larger than its critical value at the level of significance alpha=0.01. All operations were performed in R using custom routines involving the baseline package V1.3-1.

### Mating assay

Mating assays were performed as described previously^40^. Early L4 males were transferred to fresh plates and kept apart from hermaphrodites until they reached sexual maturation. Single virgin males were assayed for their mating behavior in the presence of 10–15 adult *unc-31(e928)* hermaphrodites on a plate covered with a thin fresh *E. coli* OP50 lawn. Mating behavior was scored within a 15-minute time window or until the male ejaculated, whichever occurred first. Mating was recorded using a Zeiss Axiocam ERc 5s mounted on a Zeiss stemi 508. The movie sequence was analyzed and the males were tested for their ability to perform an intact mating sequence^72^. Males were scored for their time until successful mating, contact response and vulva location efficiency. Males that did not mate within 15 minutes were not analyzed for time until successful mating and contact response parameters. Contact response requires tail apposition and initiation of backward locomotion. Percentage response to contact = 100 x (the number of times a male exhibited contact response/the number of times the male makes contact with a hermaphrodite via the rays)^40,73^. Vulva location efficiency (L.E.)^74^, or the ability to locate the vulva, was calculated as 1 divided by the number of passes or hesitations at the vulva until the male first stops at the vulva.

