## Supplemental information for "Sexually dimorphic architecture and function of a mechanosensory circuit in *C. elegans*"

### SUPPLEMNETAL INFORMATION

This file includes seven supplemental figures, a supplementary table, a supplementary file and a supplementary video.

### SUPPLEMENTAL FIGURES

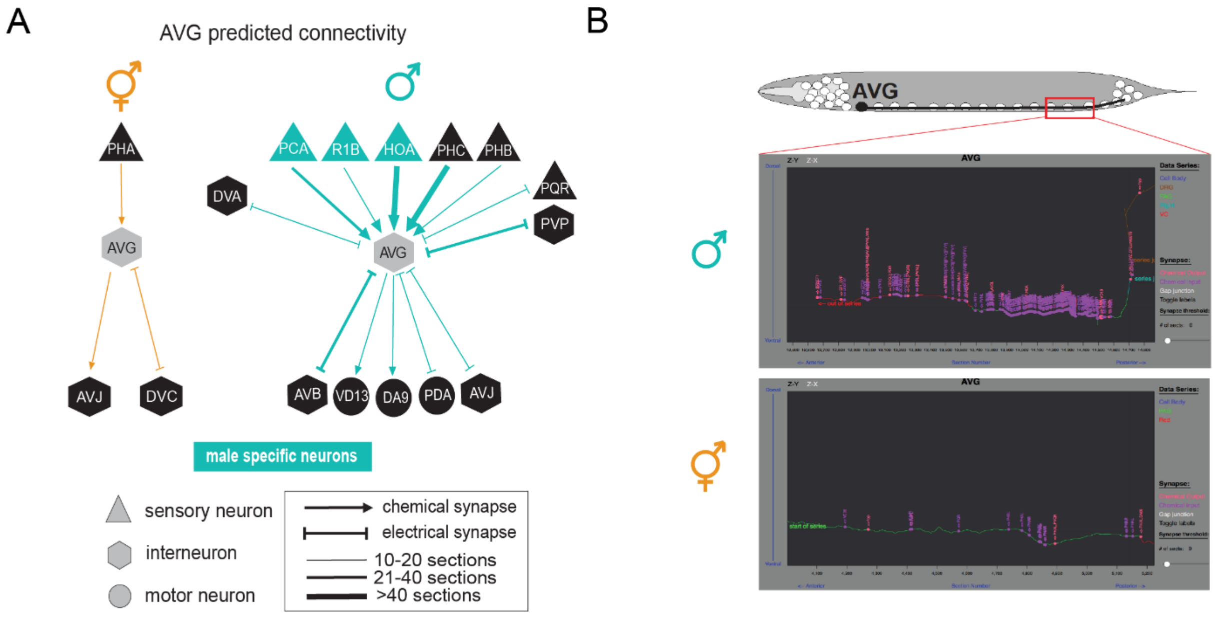


**Figure S1. The sex-shared interneuron AVG receives many inputs in males compared to hermaphrodites**

(A) Schematic diagram of the connectivity of the AVG neuron in both sexes in the adult stage based on electron microscopy reconstructions[25,27]. Chemical and electrical synapses between sensory (triangles), inter- (hexagons) and motor (circles) neurons are depicted as arrows and inhibitory arrows, accordingly. Arrow thickness correlates with the degree of connectivity (number of sections over which *en passant* synapses are observed). (B) Schematic diagram of the inputs AVG receives in males and hermaphrodites specifically in the pre-anal ganglion (PAG) in the tail of the worm (wormwiring.org). Orange- hermaphrodites, cyan- males.

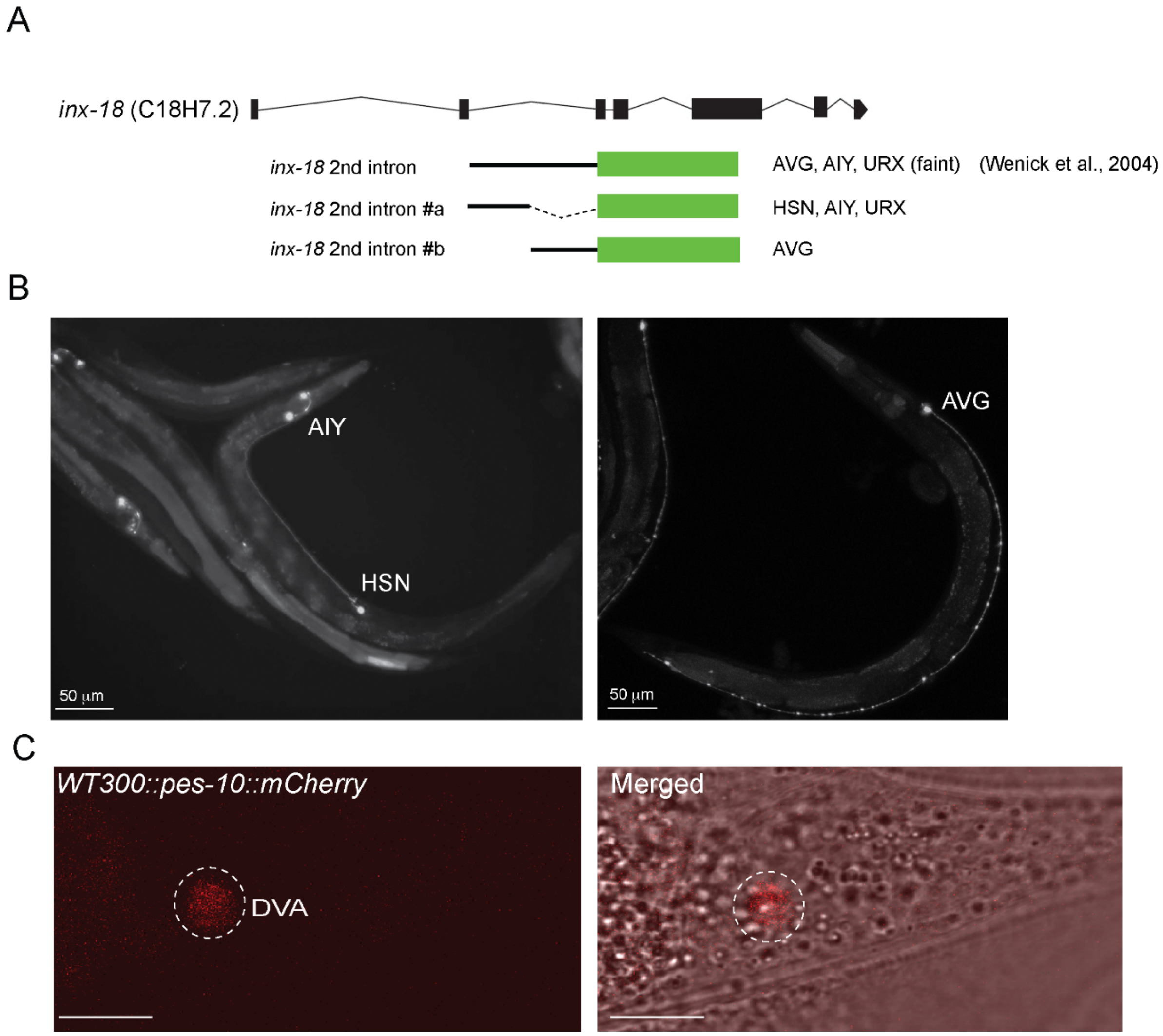


**Figure S2. Cell-specific drivers in the sex-shared interneurons AVG and DVA**

(A) Schematic of the bashing of the *inx-18* second intron to drive specific expression in AVG. Scale bar is 50 µm. (B) Representative confocal micrograph of *inx-18#a* (left) and *inx-18#b* (right). Scale bars are 50 µm. (C) Representative confocal micrograph of DVA interneuron identified by the expression of *WT300::pes-10::mCherry*[65]. Scale bars are 10 µm.

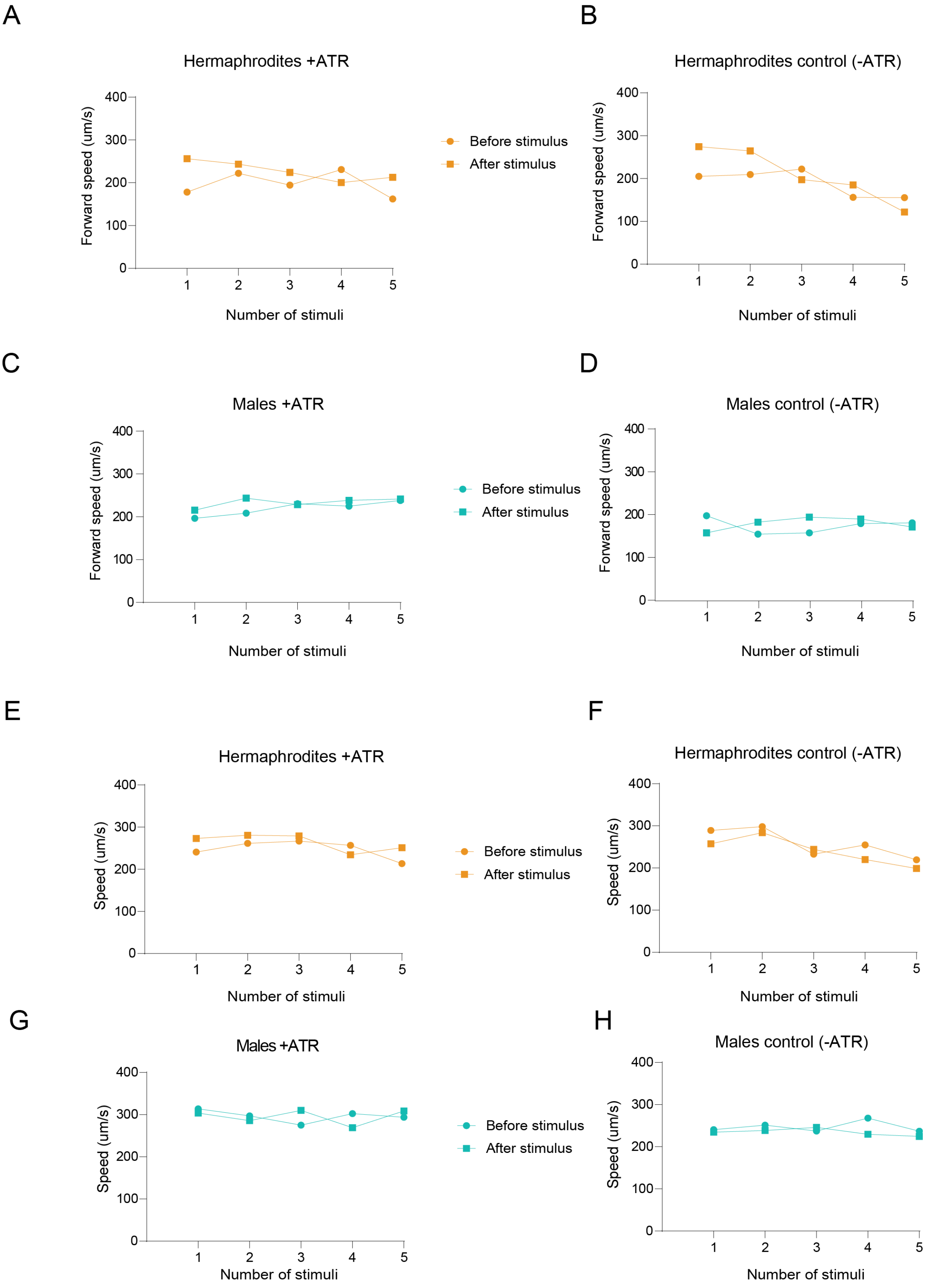


**Figure S3. Optogenetic activation of AVG does not affect the locomotion of both sexes**

The average forward speed (A-B) and total speed (forward and reverse, E-F) of hermaphrodites grown on ATR (A, E) or control (B, F) plates before and after each stimulus in a sequence of five stimuli (see *Methods*). The average forward speed (C-D) and total speed (forward and reverse, G-H) of males grown on ATR (C, G) or control (D, H) plates before and after each stimulus in a sequence of five stimuli (see *Methods*). n = 12-21 worms per group. Orange- hermaphrodites, cyan- males.

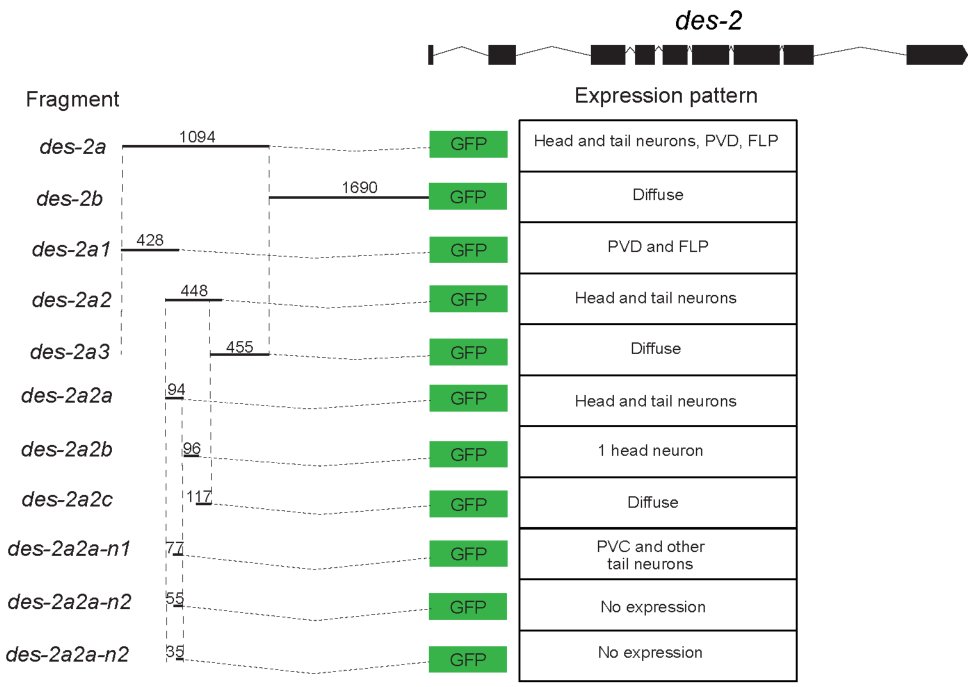


**Figure S4. *des-2* promoter analysis**

Schematic of the *des-2* promoter bashing[74]. The fragments were fused to GFP and their expression patterns were examined under confocal microscopy.

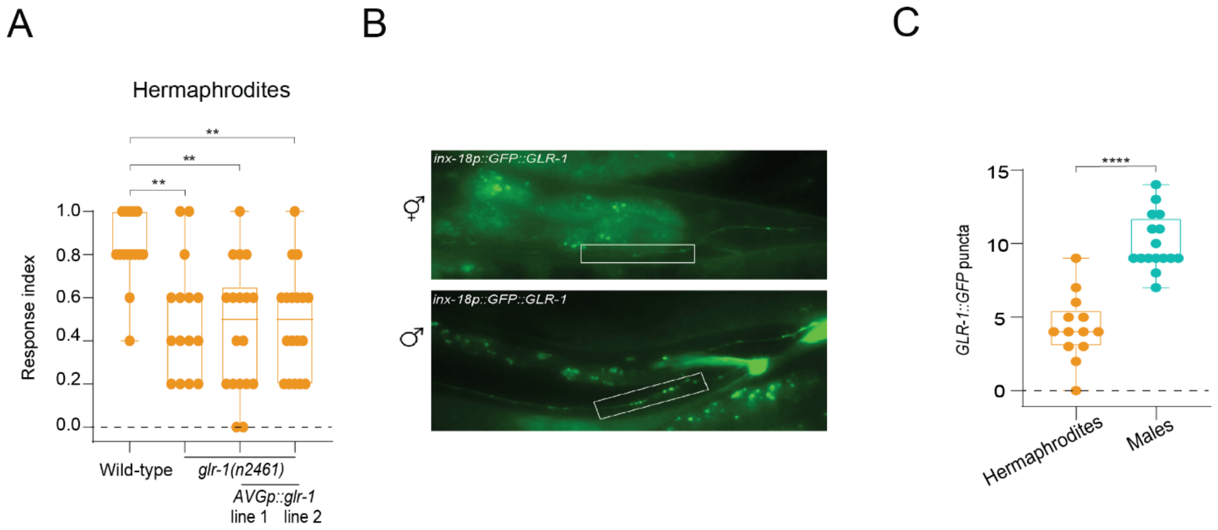


**Figure S5. *glr-1* does not function through AVG to mediate tail mechanosensation in hermaphrodites**

(A) Tail-touch responses of wild-type, *glr-1(n2461)* and two extrachromosomal lines of *glr-1(n2461);AVGp::glr-1* hermaphrodites. n=15-18 worms per group. The response index represents an average of the forward responses (scored as responded or not responded) in five assays for each animal. (B) Representative fluorescent micrographs of hermaphrodites and males expressing *inx-18p::GFP::GLR-1* in the pre-anal ganglion. White rectangles represent the area scored. (C) Quantification of GFP puncta. Worms were quantified for GFP puncta in a fixed distance of 80 µm from the anus, where most of the inputs to AVG are formed (Figure S1B). n = 13-16 worms per group. In (A) we performed a Kruskal-Wallis test followed by a Dunn's multiple comparison test, in (C) we performed Mann-Whitney test. **** $p < 0.0001$, ** $p < 0.01$. Orange- hermaphrodites, cyan- males.

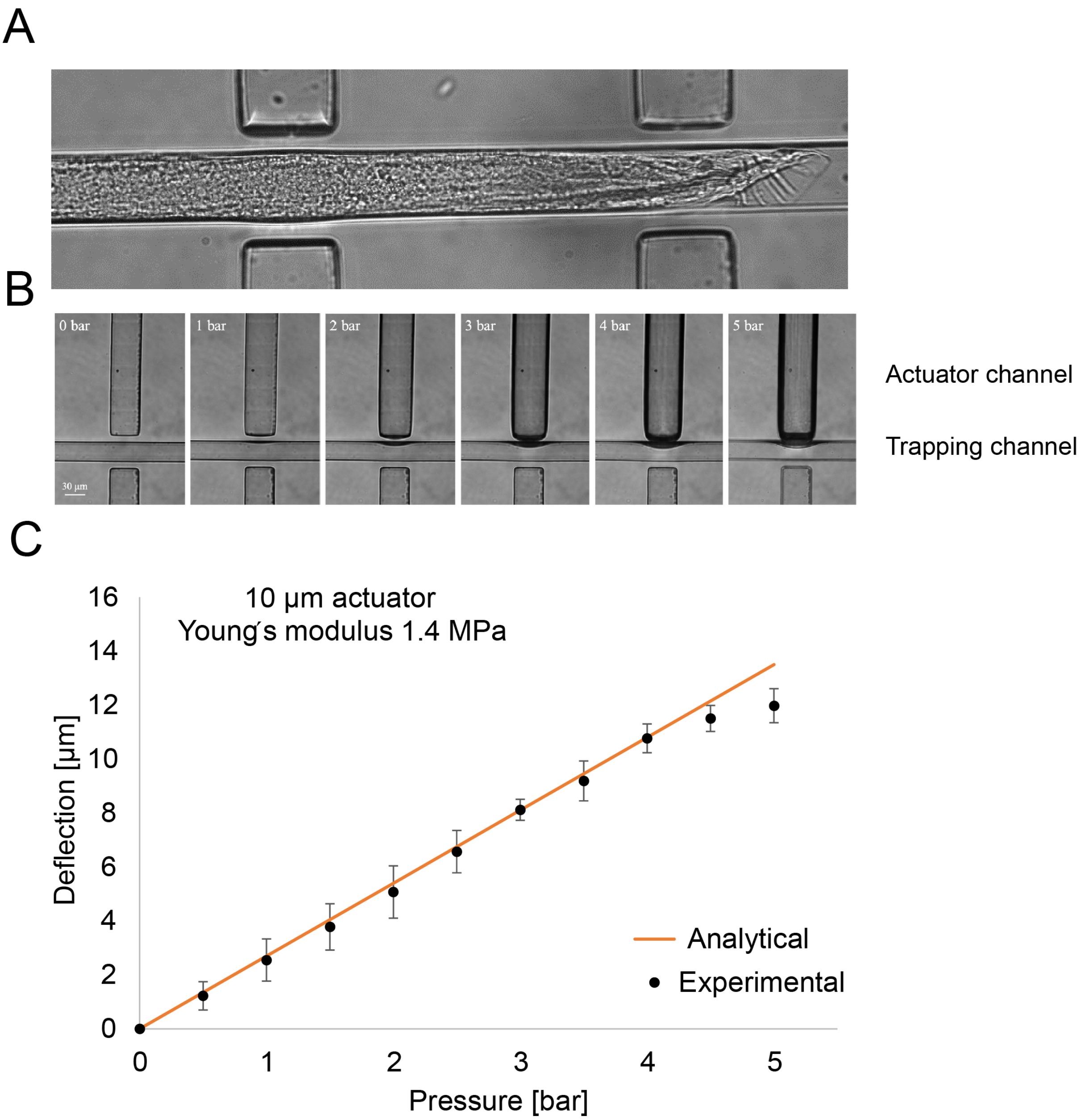


**Figure S6. Calibration of the microfluidic device designed to measure calcium activity traces in response to tail mechanical stimulation**

(A) The microscopic image of a male immobilized in a trapping channel. (B, C) Deflection of the actuator under different pressure values, from 0 to 5 bar. Deflection was measured using ImageJ without the presence of an animal inside the trapping channel.

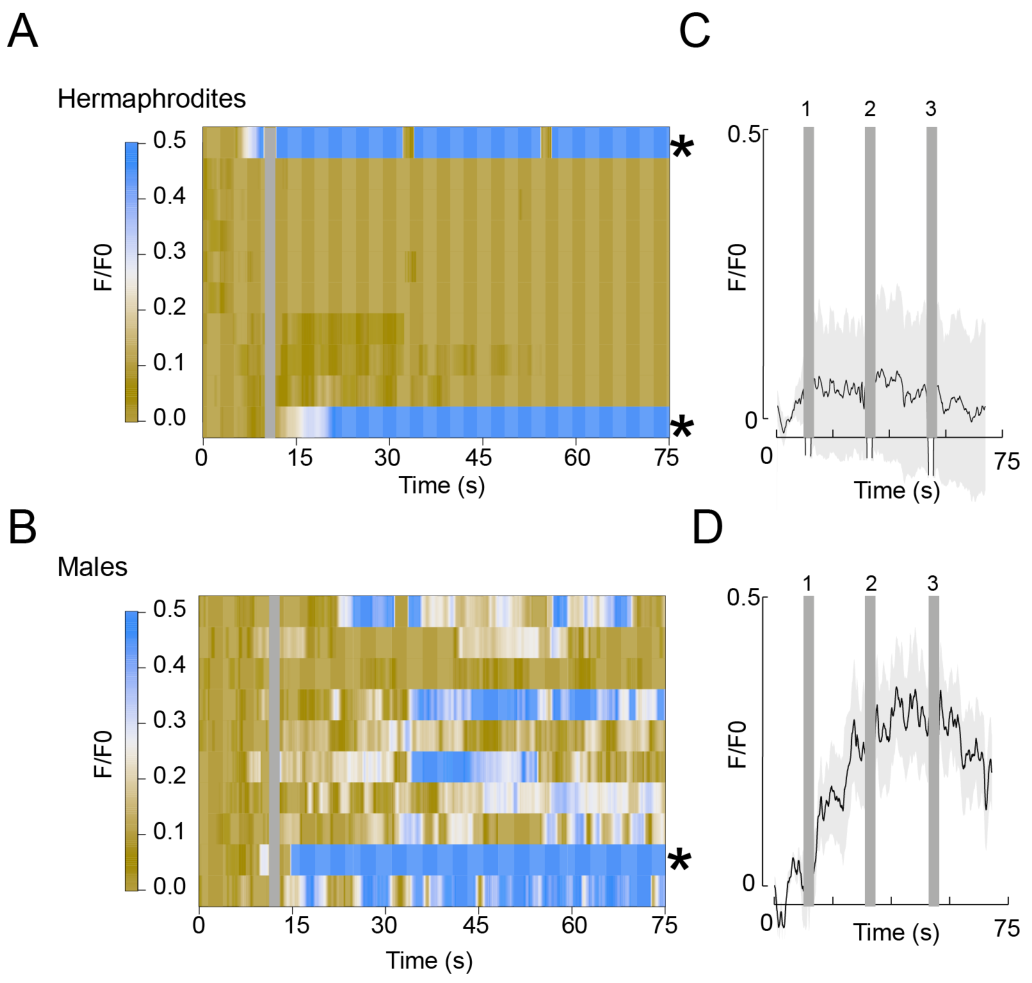


**Figure S7. AVG neuronal recordings suggest light-evoked activity**

AVG GCaMP6s calcium responses to three consecutive tail mechanical stimulations of males (A) and hermaphrodites (B). Heatmaps represent the calcium levels of individual worms. Asterisks represent individual worms where light-evoked activity is suspected to occur. Gray vertical line represents the times when the first stimulus was applied. Average and SEM traces of AVG calcium responses of males (C) and hermaphrodites (D). Gray vertical lines represent the times when a stimulus was applied. n = 9 worms per group.  Each stimulus lasted two seconds (see *Methods*).

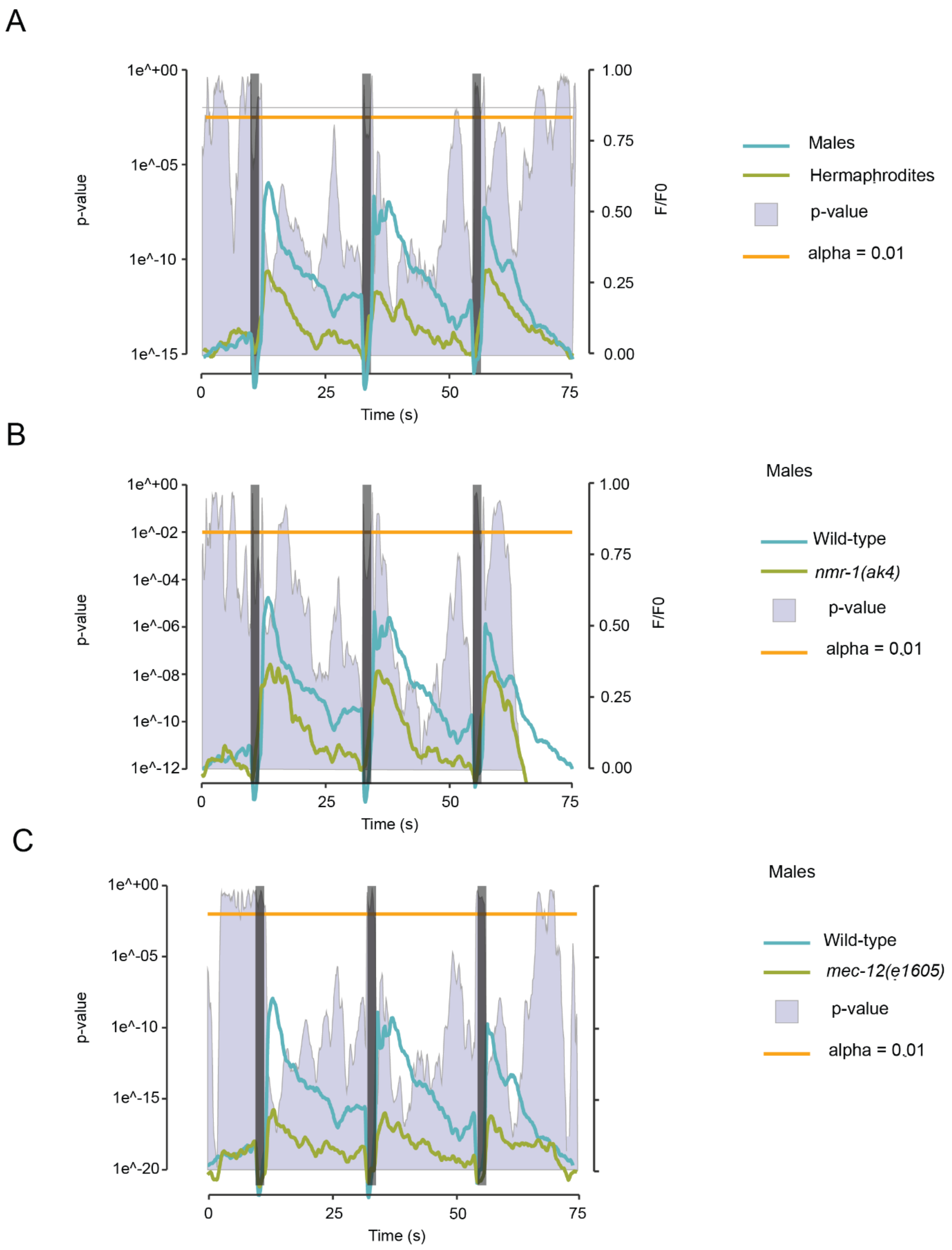


**Figure S8. Statistical representation of calcium activity traces in AVG under different conditions**

(A) The average calcium responses of males and hermaphrodites (smoothed and corrected, see *Methods*) to the running t-value in each time point (blue shaded curve). Gray vertical lines represent the times when a stimulus was applied. Orange line represents p-value = 0.01. (B, C) The average calcium responses of wild-type and *nmr-1(ak4)* males (B) and wild-type and *mec-12(e1605)* males (C) (smoothed and corrected, see *Methods*) in relation to the running t-value in each time point (blue shaded curve). Gray vertical lines represent the times when a stimulus was applied. Orange line represents p-value = 0.01.

**Supplementary Table 1. List of strains used in this study**

| Strain | Description | Source |
|---|---|---|
| CX14373 | *kyEx4571 [pNP403 (tag-HisCl1::SL2::GFP 5 ng/μL); myo-::168 [mCherry 10 ng/μL::3* | *Caenorhabditis Genetics Center* |
| OH16196 | *otEx7437[srg-13::HisCl::SL2::GFP 50ng/ul, ttx-3::mCherry 50ng/ul]; him-5 (e1490) V* | Oliver Hobert's lab |
| OH13689 | *otEx6341[gpa-6::HisCl1::SL2::GFP 50ng/ul; ttx-3:cherry 50ng/ul]; him-5(e1490) V* | *Oren-Suissa et al, 2016* |
| OH14826 | *otEx6906 [eat-4p11del11:::HisCl::SL2::GFP 50ng/μl; unc-122::gfp 50ng/μl]; him-5(e1490) V* | *Serrano-Saiz et al, 2017* |
| CB4088 | *him-5(e1490) V* | *Caenorhabditis Genetics Center* |
| CB3284 | *mec-12(e1605) III* | *Caenorhabditis Genetics Center* |
| MOS463 | *mec-12(e1605) III; him-5(e1490) V* | This study |
| RB1546 | *T13G4.3(ok1859) X* | *Caenorhabditis Genetics Center* |
| MOS478 | *T13G4.3(ok1859) X; him-5(e1490) V* | This study |
| CX4544 | *ocr-2(ak47) IV* | *Caenorhabditis Genetics Center* |
| MOS325 | *ocr-2(ak47) IV; him-5(e1490) V* | This study |
| CX10 | *osm-9(ky10) IV* | *Caenorhabditis Genetics Center* |
| MOS332 | *osm-9(ky10) IV; him-5(e1490) V* | This study |
| MOS477 | *mec-12(e1605) III; etyEx161[che-12::mec-12 30ng/ul; ttx-3::gfp 20ng/ul; pBS 50ng/ul]; him-5(e1490) V)* | This study |
| BC10751 | *dpy-5(e907) I; sEx10751* | *Caenorhabditis Genetics Center* |
| MOS481 | *dpy-5(e907) I; sEx10751; him-5(e1490) V* | This study |
| MOS505 | *etyEx174[eat-4p11::tmc-1 40ng/ul, ttx-3::gfp 30 ng/ul, pBS 30 ng/ul], T13G4.3(ok1859) X; him-5(e1490)* | This study |
| AQ4330 | *ljEx1221[Ptmc-1(4kb)::mKate2]* | *Kaulich et al, 2021* |
| OH12503 | *otIs520[eat-4p11::gfp;ttx-3::cherry]; him-5(e1490) V* | Oliver Hobert's lab |
| MOS511 | *ljEx1221[Ptmc-1(4kb)::mKate2]; otIs520(eat-4p11::gfp;ttx-3::cherry); him-5(e1490)* | This study |
| MOS182 | *etyEx188[inx-18::HisCl1::SL2::GFP 50ng/ul; ttx-3:cherry 50ng/ul]; him-5(e1490) V* | This Study |

| | | |
|---|---|---|
| MOS473 | *etyEx159[WT300::pes-10::HisCl1::SL2::GFP 50ng/ul, ttx-3::gfp 30ng/ul; pBS 20ng/u;];him-5(e1490) V* | This study |
| MOS247 | *otEx6335[inx-18::tra-2(ic)::SL2::2XNLS::tag-RFP 10ng/ul, ttx-3::GFP 50ng/ul, pBS 40ng/ul]; etyEx188[inx-18::HisCl1::SL2::GFP 50ng/ul; ttx-3:cherry 50ng/ul]; him-5(e1490)* | This study |
| MOS254 | *otIs606[inx-18p::FEM-3::SL2::wcherry; pha-1+]; etyEx188[inx-18::HisCl1::SL2::GFP 50ng/ul; ttx-3:cherry 50ng/ul]; him-5(e1490)* | This study |
| MOS195 | *etyEx44[inx-18delAIY::NpHR::mCherry 100 ng/ul + ttx-3::GFP 30 ng/ul]* | This study |
| SRS167 | *pha-1(e2123) III; lite-1(ce314) X* | Alon Zaslaver's lab |
| MOS287 | *him-5(e1490)V; lite-1(ce314) X* | This study |
| MOS203 | *etyEx44[inx-18delAIY::NpHR::mCherry 100 ng/ul + ttx-3::GFP 30 ng/ul]; him-5(e1490)* | This Study |
| KP4 | *glr-1(n2461) III* | *Caenorhabditis Genetics Center* |
| MOS340 | *glr-1(n2461) III; him-5(e1490)* | This study |
| RB1808 | *glr-2(ok2342) III* | *Caenorhabditis Genetics Center* |
| MOS326 | *glr-2(ok2342) III. ; him-5(e1490) V* | This study |
| VM487 | *nmr-1(ak4) II* | *Caenorhabditis Genetics Center* |
| MOS351 | *nmr-1(ak4) II; him-5(e1490)* | This study |
| CB1489 | *him-8(e1489) IV* | *Caenorhabditis Genetics Center* |
| VC2623 | *nmr-2(ok3324) V* | *Caenorhabditis Genetics Center* |
| MOS346 | *nmr-2(ok3324) V; him-8(e1489) IV* | This study |
| MOS434 | *etyEx137[inx-18p::nmr-1 25ng/ul; ttx-3::gfp 20 ;ng/ul; pBS 55ng/ul]; nmr-1(ak4) II; him-5(e1490) V* | This study |
| MOS70 | *otIs460 [inx-18p::wcherry; pha-1(+)]; him-8(e1489) IV* | Oliver Hobert's lab |
| MOS467 | *etyEx157[nmr-1 fosmid 10 ng/ul, ttx-3::gfp 30 ng/ul, pBS 60 ng/ul], otIs460 [inx-18p::wcherry; pha-1(+)]; him-8(e1489) IV* | This study |
| MOS239 | *otIs606 [inx-18p::FEM-3::SL2::wcherry; pha-1+]; him-5(e1490) V* | This study |

| | | |
|---|---|---|
| MOS368 | *otIs606 [inx-18p::FEM-3::SL2::wcherry; pha-1+]; nmr-1(ak4) II; him-5(e1490) V* | This study |
| MOS367 | *otIs606[inx-18p::FEM-3::SL2::wcherry; pha-1+] ;glr-1(n2461) III; him-5(e1490) V* | This study |
| MOS433 | *etyEx142[pMO32(inx-18::tra-2(ic)::SL2::2NLS) 40 ng/ul; ttx-3::gfp 30 ng/ul, pBS 30 ng/ul]* | This study |
| MOS496 | *etyEx142[pMO32(inx-18::tra-2(ic)::SL2::2NLS) 40 ng/ul; ttx-3::gfp 30 ng/ul, pBS 30 ng/ul]; him-5(e1490)* | This study |
| MOS497 | *etyEx142[pMO32(inx-18::tra-,2(ic)::SL2::2NLS) 40 ng/ul; ttx-3::gfp 30 ng/ul pBS 30 ng/ul]; glr-1(n2461)III; him-5(e1490)* | This study |
| MOS506 | *etyEx142[pMO32(inx-18::tra-,2(ic)::SL2::2NLS) 40 ng/ul; ttx-3::gfp 30 ng/ul pBS 30 ng/ul]; nmr-1(ak4) II; him-5(e1490)* | This study |
| MOS495 | *etyEx168[WT300::pes-10::nmr-1 25ng/ul, ttx-3::gfp 20 ng/ul, pBS 55 ng/ul]; nmr-1(ak4) II; him-5(e1490)* | This study |
| MOS480 | *etyEx31[inx-18b::GCaMP6s 30ng/ul, sra-6::WrmScarlet 30ng/ul, PHB::mCherry MVC15 40ng/ul]; lite-1(ce314) X; him-5(e1490)* | This study |
| MOS488 | *etyEx31[inx-18b::GCaMP6s 30ng/ul, sra-6::WrmScarlet 30ng/ul, PHB::mCherry MVC15 40ng/ul]; lite-1(ce314) X; nmr-1(ak4) II; him-5(e1490)* | This study |
| MOS511 | *etyEx31[inx-18b::GCaMP6s 30ng/ul, sra-6::WrmScarlet 30ng/ul, PHB::mCherry MVC15 40ng/ul]; lite-1(ce314) X; nmr-1(ak4) II; mec-12(e1605) III; him-5(e1490)* | This study |
| DA509 | *unc-31(e928) IV* | *Caenorhabditis Genetics Center* |
| MOS533 | *etyEx186[inx-18#b::GFP 50ng/ul, pha-1(+) 50ng/ul], pha-1(e2123)* | This study |
| MOS534 | *etyEx187[inx-18#a::GFP 50ng/ul, pha-1(+) 50ng/ul], pha-1(e2123)* | This study |
| MOS466 | *etyEx156[pHS11(WT300::pes-10::mCherry) 40ng/ul; myo-2::mCherry 5ng/ul; pBS 55ng/ul]* | This study |
| MOS214 | *etyEx54[inx-18delAIY::Chr2::mCherry 50 ng/ul + ttx-3::GFP 30 ng/ul + pBS 20 ng/ul]* | This study |
| MOS283 | *etyEx54[inx-18delAIY::Chr2::mCherry 50 ng/ul + ttx-3::GFP 30 ng/ul + pBS 20 ng/ul];him-5(e1490)V; lite-1(ce314)* | This Study |

| | | |
|---|---|---|
| MOS452 | *etyEx148(pMO41(inx-18p::GLR-1::GFP ) 40 ng/ul, ttx-3::gfp 30 ng/ul, pBS 30 ng/ul); glr-1(n2461) III; him-5(e1490) V* | This study |
| MOS453 | *etyEx149[pMO41(inx-18p::GLR-1::GFP 40ng/ul), ttx-3::gfp 30ng/ul, pBS 30ng/ul]; glr-1(n2461) III; hin-5(e1490)* | This study |
| OH13096 | *otIs582[inx-18p::GLR-1::GFP, ttx-3::mcherry]; him-5(e1490) V* | CGC |
| MOS118 | *etyEx31[inx-18b::GCaMP6s 30ng/ul, sra-6::WrmScarlet 30ng/ul, PHB::mCherry MVC15 40ng/ul]; him-5(e1490)* | *Salzberg et al, 2020* |

**Supplementary video 1**. **Representative video of calcium traces in a male AVG**

Representative video of a *lite-1*(ce314) mutant male expressing GCaMP6s in AVG, trapped inside the microfluidic device and subjected to three consecutive stimulations to the tail at the indicated time points. Only the anterior part of the animal is shown. Scale bar is 50µm. Neurons in the head are bleed-through due to the co-injection of the marker *sra-6::WrmScarlet*.

**Supplementary file 1. Autocad file with microfluidic design for mechanical stimulation to fit the size of males**
